# Heritable shell differentiation among populations of the sole lymnaeid snail across freshwaters in southern Patagonia

**DOI:** 10.64898/2026.05.14.725217

**Authors:** Micaela Müller Baigorria, Matías Abafatori, Elodie Chapuis, Nicolas Juillet, Dominique Faugère, Philippe Jarne, Patrice David, Jean-Pierre Pointier, Sylvie Hurtrez-Boussès, Pilar Alda, Nicolás Bonel

## Abstract

1. Phenotypic variation among populations in freshwater ecosystems can reflect environmentally induced responses, heritable differentiation, or both. In gastropods, shell shape varies with water flow and permanence. *Pectinidens diaphanus*, the southernmost lymnaeid worldwide, occurs in southern Patagonia, where temporary waterbodies dry out recurrently, but its geographic range and shell variation remain understudied. We reassessed its distribution and taxonomic identity, quantified reproductive and shell variation among populations from contrasting waterbodies, and evaluated whether shell differentiation reflected environmentally induced responses or persistent population differences.
2. We surveyed 190 freshwater sites between 36.2 and 54.7°S and identified *P*. *diaphanus* using morpho-anatomical examination and COI sequencing. We compared shell and reproductive variation among five populations from permanent and temporary, lentic and lotic waterbodies (a lake, a river, a stream, and two ponds). We conducted a common-garden experiment to test whether shell-shape differences between a lake and a pond population persisted after two generations.
3. *Pectinidens diaphanus* was the only lymnaeid detected in southern Patagonia. It occurred at 37 sites across four waterbody types (rivers, streams, lakes, and ponds), compared with three previously confirmed localities. Reproductive anatomy was conserved among populations, whereas shell shape differed among populations, with population origin explaining 49% of shape variation. Lake snails had more globose shells and larger apertures than stream and pond snails. Shell-shape differences between lake- and pond-derived snails persisted in the second laboratory generation but were reduced, as both lineages shifted toward more similar laboratory morphotypes, with a greater shift in the lake lineage.
4. Shell differentiation among populations was associated with water permanence and desiccation risk and persisted under common laboratory conditions. These results indicate that persistent population differences, consistent with a heritable component, coexist with environmentally induced variation in shell shape. Disentangling the contributions of genetic differentiation and phenotypic plasticity to this variation will shed light on how freshwater organisms respond to environmental heterogeneity.

## Introduction

Intraspecific variation constitutes the raw material upon which fundamental evolutionary processes such as natural selection and genetic drift act, thereby fueling diversification and adaptation (Walsh and Lynch 2018). When environmental conditions change, population persistence and subsequent evolutionary responses may depend both on the availability of heritable variation upon which selection can act and on phenotypic plasticity, which can buffer organisms against short-term environmental mismatches (Ghalambor et al. 2007; Hoffmann and Sgrò 2011). Distinguishing environmentally induced responses from heritable differentiation among populations therefore helps determine whether phenotypic differences reflect environmental plasticity or a heritable component of variation (Price et al. 2003; Pfennig 2021).

Freshwater systems provide useful contexts for studying phenotypic variation because populations experience contrasting water flow and permanence, predation pressure, and resource availability (Schluter 2000; Langerhans 2008). Morphological differentiation along environmental gradients includes variation in body shape, size, and defensive structures (Chapuis et al. 2007; Reid and Peichel 2010; Webster et al. 2011). Gastropods are particularly well suited for examining such patterns, as shell architecture exhibits variability within and among species and provides a model for evolutionary biology (Vermeij 1987; Samadi et al. 2000). Shell shape can vary continuously within species, even over small spatial scales (Pfenninger et al. 2006; Correa et al. 2011). Differences in shell thickness, aperture size, and spire height are linked to desiccation risk, hydrodynamic stress, and predation, reflecting the potential roles of environmental selection, phenotypic plasticity, and genetic differentiation (Trussell 2000, Bourdeau et al. 2015, Bonel et al. 2021). Two of these conditions, water flow and permanence, can vary independently, as both lotic and lentic waterbodies may be permanent or temporary. Populations from lotic waterbodies tend to have larger apertures than those from lentic waterbodies (Lam and Calow 1988; Facon et al. 2004; Poznańska et al. 2015). In contrast, populations from temporary waterbodies generally have more elongated shells and smaller apertures than those from permanent ones, a pattern attributed to selection against desiccation under recurrent waterbody contraction and periodic drying (Bourdeau et al. 2015). Shell phenotypes can also change under common-garden conditions and respond to selection (Gustafson et al. 2014; Noël et al. 2017). Shell differentiation among populations can therefore reflect both environmental conditions and persistent population differences.

Shell variation can also complicate species delimitation in gastropods because similar environmental conditions may produce convergent forms, whereas substantial within-species variation may be mistaken for taxonomic differentiation (Trussell 2000; Alda et al. 2021; Whelan 2021). These challenges are well documented in lymnaeid snails, a group in which shell morphology can be highly variable and may provide insufficient diagnostic resolution on its own. Integrative taxonomic approaches combine shell morphology with molecular and anatomical evidence (Padial et al. 2010). In lymnaeids, reproductive structures can provide useful diagnostic information where shell morphology is variable (Vinarski and Glöer 2009; Vinarski and Pointier 2023). Accurate species identification is relevant to understanding the transmission of trematode infections, as several lymnaeid species act as intermediate hosts of *Fasciola hepatica*, the causative agent of fasciolosis (Vázquez et al. 2023). In southern Patagonia, where fasciolosis is highly prevalent and occurs in livestock (Aguilar and Olaechea 2014; Larroza et al. 2023), determining the distribution of local lymnaeid species can help identify areas where potential intermediate hosts occur.

The South American species *Pectinidens diaphanus* is of particular ecological and biogeographical interest, as it is the southernmost-distributed lymnaeid worldwide and the only representative of the family in southern Patagonia. However, *P*. *diaphanus* remains understudied in terms of its geographic range and the basis of shell variation among populations. Historical records based on shell morphology suggested a wide American distribution, but morphological, anatomical, and molecular studies revealed that many records were misidentifications (Baker 1911; Correa et al. 2010; Vinarski and Pointier 2023). Subsequent analyses restricted its distribution to three localities in southern Patagonia (Fig. 1A; Duffy et al. 2009; Correa et al. 2010; Bargues et al. 2012). The extent to which these records represent the species’ geographic distribution across southern Patagonia remains to be established. Moreover, the relative contributions of environmental responses and persistent population differentiation to shell variation among populations from contrasting freshwaters remain unresolved. Common-garden experiments can be used to test whether population differences persist under standardized conditions. If phenotypic differentiation is primarily environmentally induced, differences between populations should decrease under common conditions, whereas persistent differences would support a heritable component (Lam and Calow 1988; Gustafson et al. 2014).

**Figure 1.**
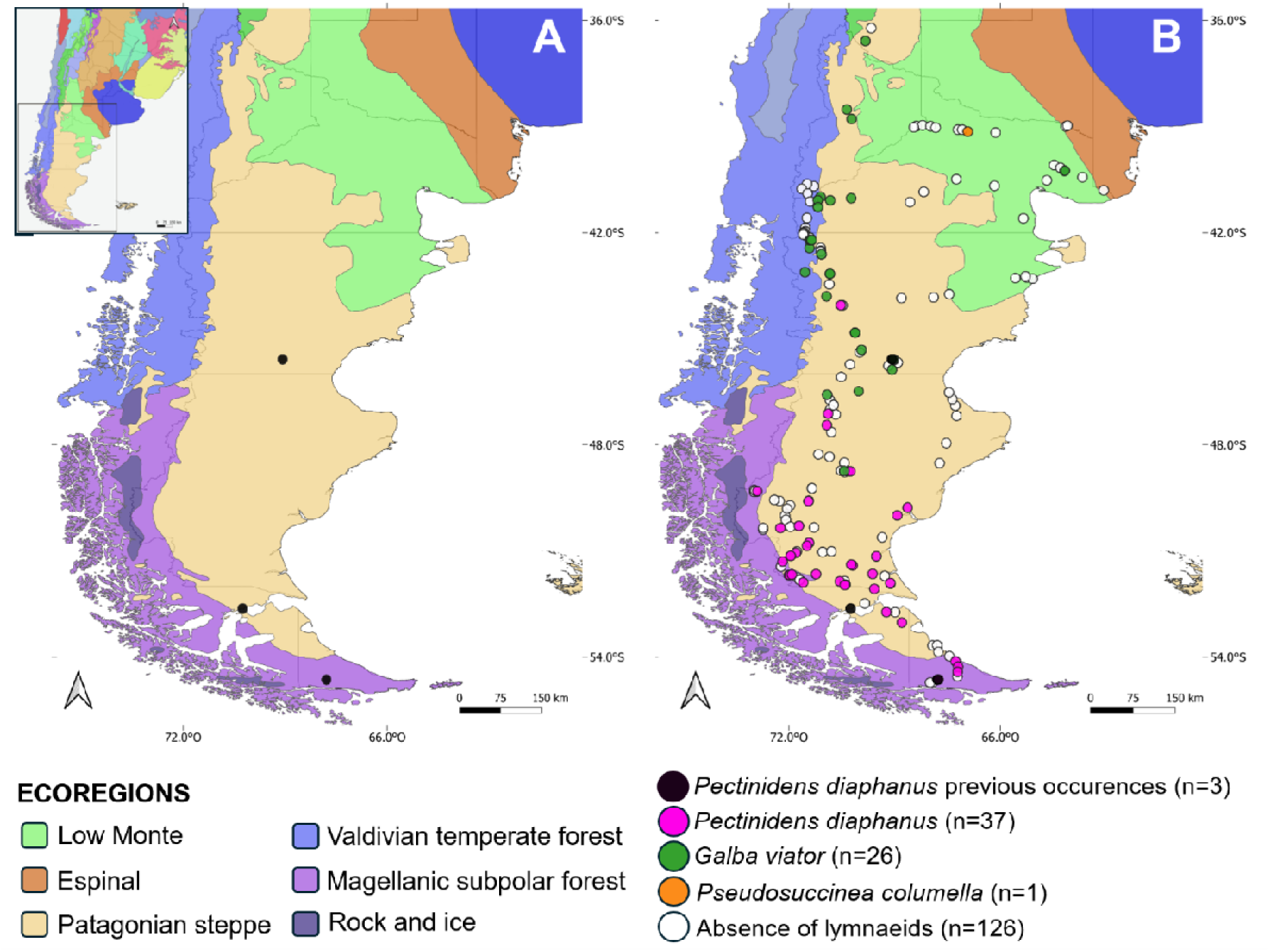
Distribution of *Pectinidens diaphanus* records in Patagonia. (A) Previously published occurrences of *P*. *diaphanus* based on Duffy et al. (2009); Correa et al. (2010); Bargues et al. (2012). Ecoregion boundaries obtained from Olson et al. (2001). (B) Distribution of lymnaeid sampling sites and *P. diaphanus* records based on field surveys. Other lymnaeid species recorded include *Galba viator* and *Pseudosuccinea columella*. Geographic coordinates of all sampling sites are provided in Table S1.

Our study had three objectives. First, we reassessed the distribution and taxonomic identity of *P. diaphanus* across southern Patagonia using field surveys combined with morphological, anatomical, and molecular evidence. Second, we quantified reproductive anatomy and shell shape among populations occupying contrasting freshwater ecosystems. Based on previous studies, we predict that populations from temporary waterbodies would exhibit more elongated shells and relatively smaller apertures than populations from permanent waterbodies (H_1_). Third, we tested whether shell differentiation between a permanent lake population and a temporary pond population persisted after two generations under common-garden conditions. We predict that shell shape differences would persist, at least in part, under common-garden conditions if differentiation includes a heritable component, whereas differences driven primarily by environmental effects would diminish (H_2_).

## Materials and methods

### Assessing geographic range and taxonomic identity

#### Field methods

To evaluate the distribution of the freshwater snail *Pectinidens diaphanus*, we conducted seven field campaigns between 2004 and 2026 in southern Argentina and Chile (southern Patagonia), spanning approximately 18.5° of latitude (36.2–54.7°S) and 10° of longitude (63.1–73.0°W) (Table S1), visiting 190 sites in total. Eight sites were revisited during the subsequent sampling campaign: Chorrillo de los Frailes, Cancha Carrera, Las Horquetas, Laguna Azul, Potrok Aike, Potrok Aike source, Frías, and Isla Pavón. We surveyed a wide range of freshwater waterbodies, including lakes (n=17), rivers (n=80), streams (n=65), ponds (n=9), inflows (n=5), flooded meadows (n=7), dry ditches (n=2), water trough (n=2), water tank (n=1), water tank + trough (n=1), stream + pond (n=1). We sampled along the shoreline by foraging at depths of up to *ca*. 80 cm. We collected snails using a strainer in soft-bottomed waterbodies and by hand in rocky substrates. We recorded GPS coordinates and assessed the presence or absence of lymnaeid snails. Surveys were standardized in duration and conducted by at least two persons. Because sampling consisted of single-visit surveys at most sites, non-detection should not necessarily be interpreted as evidence of species absence, particularly where snail abundance was low. Accordingly, our distribution assessment was based primarily on confirmed occurrences rather than on the absence of records at surveyed sites. In the field, we distinguished lymnaeids from other freshwater gastropods, primarily *Physa acuta* (sinistral shell orientation) and *Chilina* spp. (characteristic vertical brownish wavy color patterns; Cuezzo et al. 2020), but we did not identify them to species level. When we detected lymnaeids, we manually collected 10-30 individuals per site and immersed them in water at 70 °C for 30–45 s (Alda et al. 2021). We then removed soft tissues from the shell and preserved both tissues and shells in 70% ethanol for anatomical, morphometric, and molecular analyses.

#### From species identification to a distribution map

To distinguish *P. diaphanus* from the other lymnaeid snail present in Patagonia, *Galba viator*, we used an integrative approach combining detailed morpho-anatomical assessments with molecular analyses (Pointier et al. 2009; Bargues et al. 2012). Morpho-anatomical identification was based on external and internal anatomical features widely used in lymnaeid taxonomy (Pointier et al. 2009). We examined shell shape and internal anatomy of 10 individuals per site following Pointier and Vázquez (2020). Key diagnostic characters, particularly reproductive anatomy, are only fully developed in mature specimens. Therefore, species identification was based primarily on adult individuals (shell length: 10–15 mm) to avoid misclassifying undeveloped juveniles, based on observation under a stereomicroscope. Shells showed shallow sutures and lacked marked periostracal sculpture. The reproductive system featured a sinuous spermiduct, a straight ureter, a penis sheath equal in length to the preputium, and a prostate with a longitudinal fissure (Pointier and Vázquez 2020).

For the molecular analysis, we extracted DNA from one or two individuals per population morphologically identified as *P*. *diaphanus*. DNA extraction, amplification of the cytochrome *c* oxidase subunit I (COI) gene, and sequencing were carried out following Alda et al. (2021). Briefly, approximately one-third of the foot tissue was dissected from each individual, and excess ethanol was removed using sterile paper towels. Genomic DNA was then extracted using a 5% Chelex® solution (Bio-Rad) according to the manufacturer’s instructions. We amplified a fragment of the COI gene (473–627 pb) using primers LCO1490 and HCO2198 (Folmer et al. 1994). We conducted PCR using Qiagen PCR Master Mix, following manufacturer’s instruction. We visualized PCR products on a 2% agarose gel stained with EZ-Vision DNA dye and compared against a 100 bp molecular ladder. Positive amplicons were purified and sequenced by Sanger sequencing (Eurofins Genomics, Ebersberg, Germany).

We deposited the resulting sequences in GenBank (accession numbers: PZ427721–PZ427760) and compared them using BLAST with three reference sequences of *P. diaphanus* (GenBank: JF909501, Bargues et al. 2012; JN614399 and JN614400, Correa et al. 2010). To further assess species identification, we examined the relationships between newly generated sequences and published *P. diaphanus* reference sequences using a haplotype network, including *Ladislavella elodes* (GenBank: MF544168) as an outgroup, the phylogenetically closest species to *P. diaphanus* (Correa et al. 2010). We aligned sequences using MAFFT with default parameters (Katoh and Standley 2013), and the alignment was trimmed using Gblocks with less stringent settings. We built and visualized a haplotype network in PopART v4.8.4 (Leigh and Bryant 2015) using the statistical parsimony method implemented in TCS (Clement et al. 2002).

We compiled spatial occurrence data using sampling coordinates to produce a distribution map of *P*. *diaphanus* across Patagonia. We performed mapping using QGIS (version 3.30.0) and classified each site by the presence of *P*. *diaphanus*, other lymnaeid species, and the absence of lymnaeids.

### Quantifying reproductive and shell morphological variation among populations

To investigate reproductive and shell variation, we sampled five populations of *P*. *diaphanus* in November 2023, representing four waterbody categories: one lake, one river, one stream, and two ponds (Table S2; Fig. S1). The sampling sites were located within a restricted geographic area in southern Santa Cruz Province, Argentina, spanning approximately 1° of latitude (51.1–52.1°S), with pairwise distances ranging from approximately 8 to 200 km (Fig. S1). This geographic proximity helped minimize regional climatic differences among sampling sites We transported 80–100 live snails from each site to the laboratory and maintained them in two separate 1.5-L aquaria per population containing spring water and continuous aeration (G_0_ generation). Snails were kept under controlled conditions at 16 ± 1 °C, with a 16:8 h light/dark cycle, and fed ad libitum with boiled and ground lettuce.

We classified waterbody types using broad operational hydrological and structural criteria rather than strict geomorphological definitions. We defined lentic waters as waterbodies with no perceptible surface flow (e.g. lakes and ponds), and lotic waters as waterbodies with perceptible surface flow (e.g. rivers and streams; Wetzel 2001; Davies et al. 2004). We generally classified ponds as shallow (maximum depth < 2 m) and small (surface area < 1 ha), and lakes as deeper (> 2 m) and larger (> 1 ha). Similarly, we classified streams primarily by their wetted channel width (< 15 m), which was generally accompanied by shallow water (mean depth < 0.5 m), whereas we classified rivers by their wider channels (> 15 m), greater depths, and continuous baseflow. We used these structural criteria as general guidelines rather than strict thresholds and also considered our field observations in this characterization.

Continuous hydrological measurements were not available for the sampled sites. We classified waterbody types and their associated permanence regimes based on our knowledge of the sites and field observations accumulated during sampling campaigns and revisits between 2004 and 2026. Where available, we also consulted local hydrological records and considered visible evidence of seasonal desiccation (e.g. dry channel margins or desiccated basins) in the context of regional climate (Mayr et al. 2007; Fovet et al. 2021). These classifications represent broad, qualitative contrasts in water flow and permanence (lentic vs. lotic; perennial vs. intermittent/ephemeral), rather than site-specific estimates based on continuous hydrological monitoring.

#### Morphometry of the reproductive system in wild populations

We dissected ten adult snails from each of the five selected populations under a stereomicroscope. To minimize potential age-related variation, we selected individuals with four shell whorls, whorl number being a proxy for age (Vermeij 1973; Kemp and Bertness 1984). We dissected and illustrated the prostate and penial complex using a camera lucida attached to a stereomicroscope. Illustrations were digitized using a flatbed scanner, and reproductive traits (area and perimeter) of both the prostate and penial complex were quantified using *ImageJ* software (Schneider et al. 2012). We evaluated differences in reproductive traits among populations using linear models (LMs), with population origin as a fixed effect and four reproductive traits as response variables. Error distributions were modeled using a Gaussian distribution, and model assumptions, including normality and homoscedasticity of residuals, were checked by visual inspection of diagnostic plots. Analyses were conducted in *R* v4.3.1 (R Core Team 2024) using the base statistical functions implemented in the *stats* package.

#### Shell morphometrics in wild populations

To assess variation in shell shape, we performed geometric morphometric analyses on adult individuals from the same five *P. diaphanus* populations used for reproductive morphology analyses. We took 107 high-resolution shell photographs (10–22 individuals per population) (Albarrán-Mélzer et al. 2020). We positioned specimens in apertural view and digitized images using *tpsUtil* v1.82 (Rohlf 2015). Landmark-based morphometric analysis was carried out in *tpsDig2* v2.31 (Rohlf 2015), using 15 homologous landmarks per shell (Fig. 2; e.g. Tamburi et al. 2018), including points placed along the aperture margin at consistent anatomical positions. We digitized landmark configurations and initially aligned them using Generalized Procrustes Analysis (GPA) in *MorphoJ* v1.08.02 (Klingenberg 2011). We conducted subsequent analyses in *R* using the package *geomorph* v4.0.10 (Adams and Otárola-Castillo 2013), where centroid size (CS) was extracted and Procrustes coordinates were used for multivariate analyses of shape variation. To evaluate allometric effects, shape coordinates were regressed on CS using a permutation-based Procrustes analysis of variance (Procrustes ANOVA) implemented with the *procD.lm* function in *geomorph*. Because CS had a small but significant effect on shell shape, it was included as a covariate in subsequent analyses to account for allometry.

**Figure 2.**
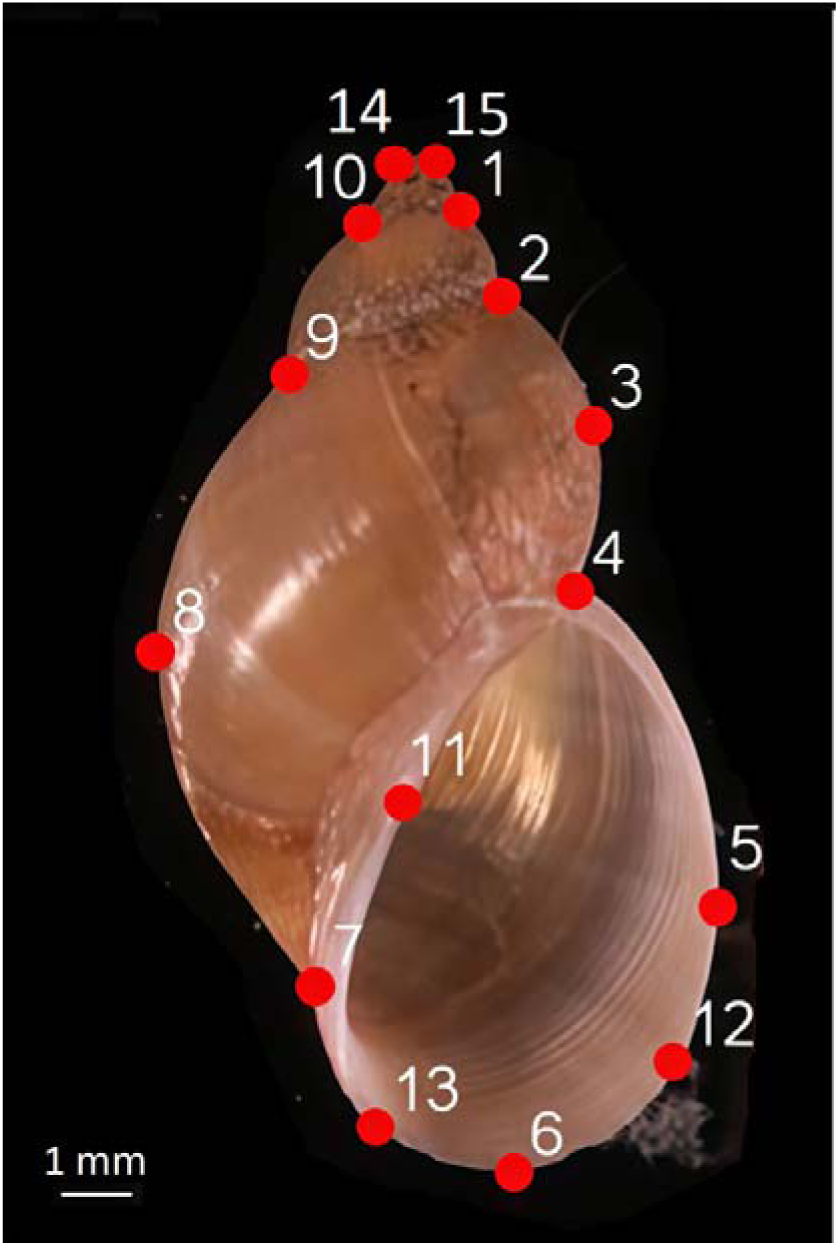
Landmark configuration used for geometric morphometric analyses of *Pectinidens diaphanus* shells. Fifteen homologous landmarks were digitized on each shell, including points distributed along the aperture margin and key anatomical regions of the shell outline. Landmark positions are shown on a representative specimen. The same configuration was applied to wild individuals (G_0_) and second-generation laboratory-reared individuals (G_2_).

We first explored shape variation using Principal Component Analysis (PCA) based on Procrustes coordinates. Next, we tested differences in shell shape among populations using Procrustes ANOVA, with significance assessed through 10,000 permutations. When the global test was significant, pairwise comparisons with Holm-Bonferroni adjustment between populations were performed using the same framework. To examine multivariate differentiation among wild populations, we performed a Canonical Variate Analysis (CVA) on Procrustes coordinates. We used CVA as an exploratory tool to visualize population separation in morphospace, which maximizes among-population relative to within-population variation. Differences in mean shape between selected population pairs were visualized using thin-plate spline (TPS) deformation grids derived from landmark configurations, illustrating landmark displacements associated with morphological variation.

### Common-garden experiment and assessment of heritable shell shape variation

To evaluate whether shell shape differences in wild populations (G_0_) persisted under common laboratory conditions, we randomly selected snails from a lake and a pond population known to exhibit contrasting shell morphologies. We maintained field-collected individuals under the laboratory conditions described above and bred them to the second laboratory generation (G_2_) in population-specific breeding lines through mass mating (Fig. S2). Rearing snails for two generations under common conditions allowed us to minimize maternal and environmental carry-over effects associated with wild-caught parents and their offspring (Falconer and Mackay 1996; Mousseau and Fox 1998). Because individuals were bred through mass mating rather than in controlled family groups, this design did not allow us to partition phenotypic variance into genetic and environmental components or estimate heritability. Instead, it allowed us to assess whether shell shape differences between wild populations persisted across generations under standardized environmental conditions. We analyzed shell shape variation in the G_2_ generation (N = 33 individuals from the lake, N = 39 from the pond) using geometric morphometric methods based on landmark coordinates as described for G_0_ snails.

To test whether laboratory conditions altered shell shape differently between source populations, we analyzed the lake and pond populations represented in both generations (G_0_: lake, n = 39; pond, n = 20; G_2_: lake-derived, n = 33; pond-derived, n = 39). Only the Chorrillo de los Frailes pond sample was used for G_0_, as this is the source of the G_2_ pond line. We fitted a Procrustes ANOVA with log-transformed centroid size (logCS, mm), population origin (lake vs. pond), generation (G_0_ vs. G_2_) and their interaction, using *procD.lm* in *geomorph*. Terms were tested sequentially (type I sums of squares) in that order: logCS first to account for allometry, then population to capture pre-existing differentiation, generation to capture the effect of laboratory rearing, and their interaction to test whether that effect differed between populations. Significance was assessed by residual randomization (10,000 permutations). Differences among the four population × generation groups were tested with pairwise Procrustes ANOVAs (shape ∼ group + logCS), with *P* values adjusted by the Holm-Bonferroni method. To visualize rearing-induced shifts relative to population differentiation, we performed an exploratory CVA on the combined G_0_–G_2_ dataset using six groups (stream, river, pond G_0_, lake G_0_, pond G_2_, and lake G_2_). Finally, we compared the morphological trajectories of the two populations using a PTA (Adams and Collyer 2009; Adams and Otárola-Castillo 2013; *trajectory.analysis* in *geomorph*), fitted to the same model. Each trajectory was defined as the vector connecting the mean G_0_ shape to the mean G_2_ shape of a population. We compared trajectory length (path distance), the absolute difference in length between populations, and trajectory direction (angle between vectors, from the vector correlation) with 10,000 permutations. Trajectories were projected onto the first two principal components of fitted shape for visualization.

## Results

### Geographical range and species occurrence

We recorded freshwater snails at 108 of 190 sites (57%) including lymnaeids at 64 sites (34%). Chilinid snails occurred at 50 sites, cohabiting with lymnaeids at 17 of them. *Galba viator* was recorded at 26 sites in the northern survey area (Buenos Aires, Neuquén, Río Negro, Chubut, and Santa Cruz provinces, Argentina; Fig. 1B), whereas another lymnaeid species, *Pseudosuccinea columella*, was observed at a single locality in Río Negro province (Fig. 1B). In contrast, *P. diaphanus* occurred at 37 sites across southern Patagonia, including Santa Cruz Province in Argentina and the island of Tierra del Fuego in Argentina and Chile. These sites spanned four waterbody types: rivers (n = 20), streams (n = 11), lakes (n = 3), and ponds (n = 3), with most populations (84%) occurring in the Patagonian steppe ecoregion (Fig. 1B). Individuals were identified as *P*. *diaphanus* based on shell traits and reproductive anatomy (Fig. S3). Molecular analyses further supported this identification: all sequences showed 97–99% identity to reference sequences of *P. diaphanus* and the TCS haplotype network grouped newly generated sequences together with previously published references, separated by 70 mutations from the closest relative *Ladislavella elodes* (Fig. S4). No other lymnaeid species were detected at sites where *P*. *diaphanus* occurred.

### Reproductive morphology and shell shape variation among wild populations

Reproductive anatomical traits were similar among populations (Table S3). Although overall ANOVA models detected variation in prostate perimeter and penial complex dimensions, pairwise comparisons revealed no significant differences except for penial complex perimeter between river and stream populations (Table S4).

Shell shape differed markedly among wild populations (Procrustes ANOVA, *F*_3, 102_ = 34.53, *P* <0.001), with population origin explaining 49.3% of total shape variation. Centroid size had a significant effect on shell shape (*F*_1, 102_ = 4.67, *P* <0.001), but its contribution was minor compared to population origin (partial R^2^ = 0.022). All pairs of populations differed significantly in shell shape (pairwise Procrustes ANOVA, *P_Holm_*< 0.005 in all cases; Table S5), with effect sizes (R^2^) ranging from 6.8% (stream vs. pond) to 49.6% (pond vs. lake). The separation among populations in morphospace was captured by three canonical variate axes, of which CV1 and CV2 accounted for 75.8% and 13.2% of the variation, respectively (CVA; Fig. 3A). Lake and river populations formed distinct clusters, whereas stream and pond populations occupied neighboring regions of morphospace. Lake individuals had more globose shells with larger apertures, whereas stream and pond individuals had more elongated shells and smaller apertures (TPS; Fig. 3B).

**Figure 3.**
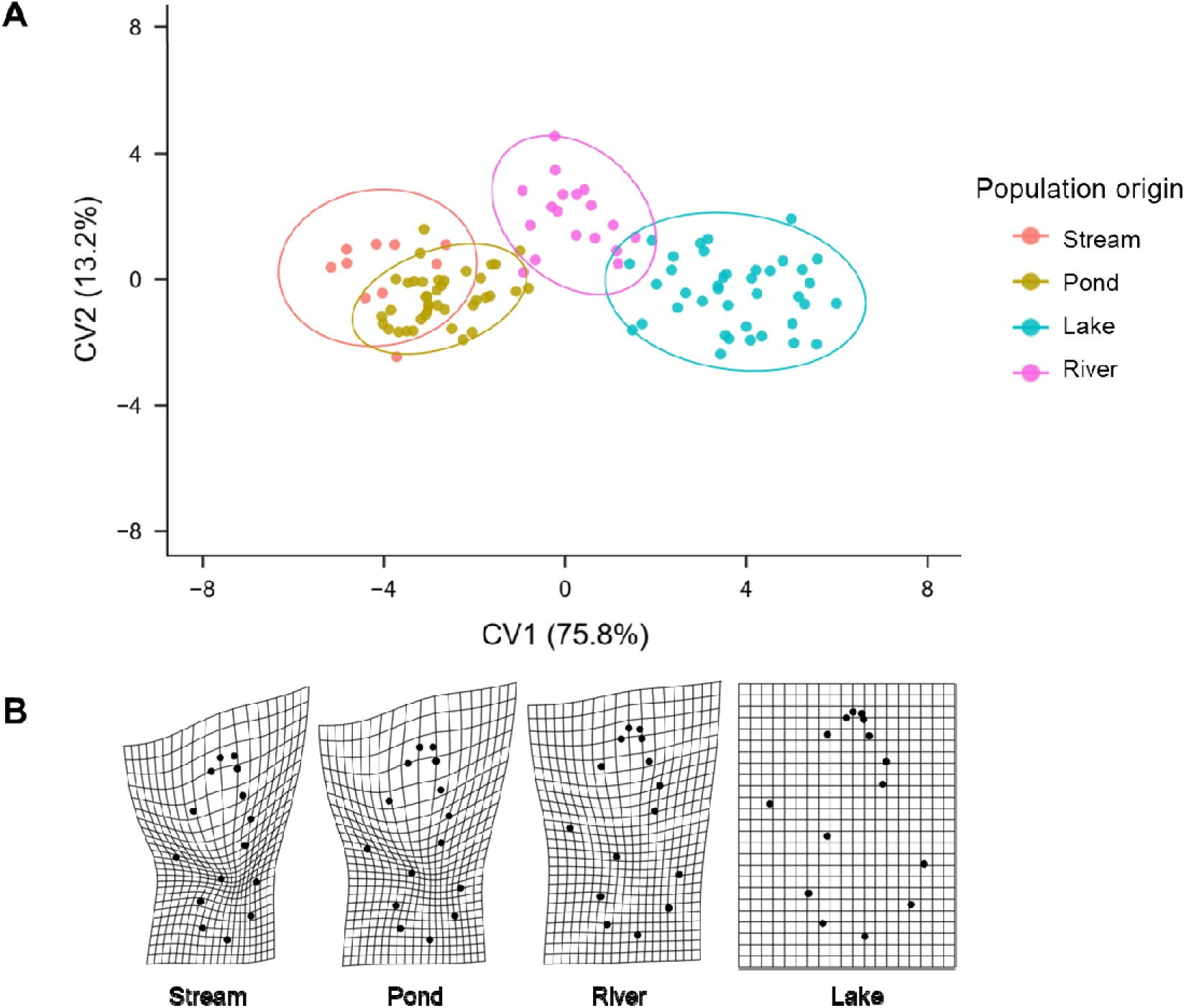
Shell shape differentiation among natural populations of the lymnaeid snail *Pectinidens diaphanus*. (A) Canonical variate analysis based on 15 homologous landmarks from individuals collected across four waterbody types (lake, river, stream, and pond). Scatterplot of individual scores along first and second canonical variates (CV1 and CV2) derived from Procrustes-aligned coordinates. Points represent individual shells, colored according to population of origin. Ellipses represent 95% data for each population. (B) Thin-plate spline deformation grids illustrating differences in mean shell shape among populations, with the lake mean shape used as reference.

### Shell shape variation under common garden conditions

In G_2_, shell shape also differed significantly between pond- and lake-derived individuals after accounting for centroid size (Procrustes ANOVA, R^2^ = 0.104, *F*_1, 69_ = 8.85, *P* < 0.001). Centroid size also explained a partial effect (partial R^2^ = 0.087, *F*_1, 69_ = 7.46, *P* < 0.001). Pond- and lake-derived G_2_ individuals occupied distinct regions of morphospace along CV1, accounting for 100% of the between-group variation (CVA; Fig. 4A). Pond-derived G_2_ individuals exhibited elongated shells, whereas lake-derived individuals displayed more globose morphologies (TPS; Fig. 4B).

**Figure 4.**
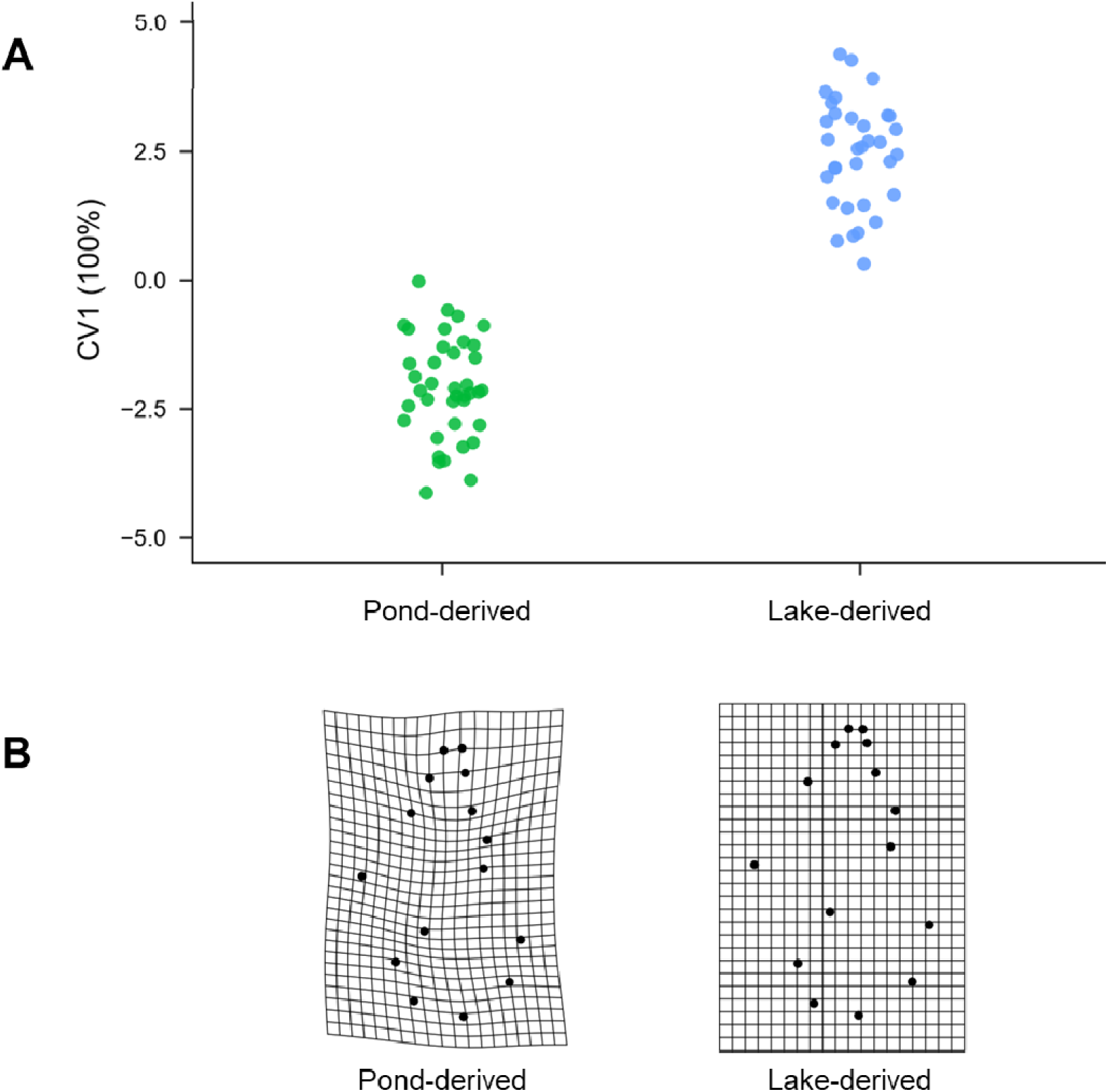
Shell shape differentiation under common-garden conditions in *Pectinidens diaphanus*. (A) Canonical variate analysis of G_2_ individuals derived from lake and pond populations and reared under standardized laboratory conditions. Because only two groups were compared, a single canonical axis (CV1) was obtained, accounting for 100% of the between-group variation. Points represent individual shells, colored according to population of origin. (B) Thin-plate spline deformation grids illustrating differences in mean shell shape between lake- and pond-derived G_2_ individuals (see Fig. 3 for details).

### Morphological changes across generations

Shell shape differed significantly among the six populations representing wild (G_0_) and laboratory-reared (G_2_) snails (Procrustes ANOVA, R^2^ = 0.468, *F*_5, 172_ = 31.21, *P* < 0.001; Table S6A). Centroid size also had a significant effect on shell shape (R^2^ = 0.017, *F*_1, 172_ = 5.55, *P* < 0.001). All pairwise population comparisons were significant (*P*_Holm_ < 0.005 in all cases; Table S6B). Wild (G_0_) and laboratory-reared (G_2_) individuals occupied different regions of morphospace; the CVA yielded five canonical axes, with CV1 and CV2 for 52.9% and 19.6% of the variation, respectively (Fig. 5A). Among G_2_ individuals, lake- and pond-derived snails still occupied different regions of morphospace, although some overlap remained.

**Figure 5.**
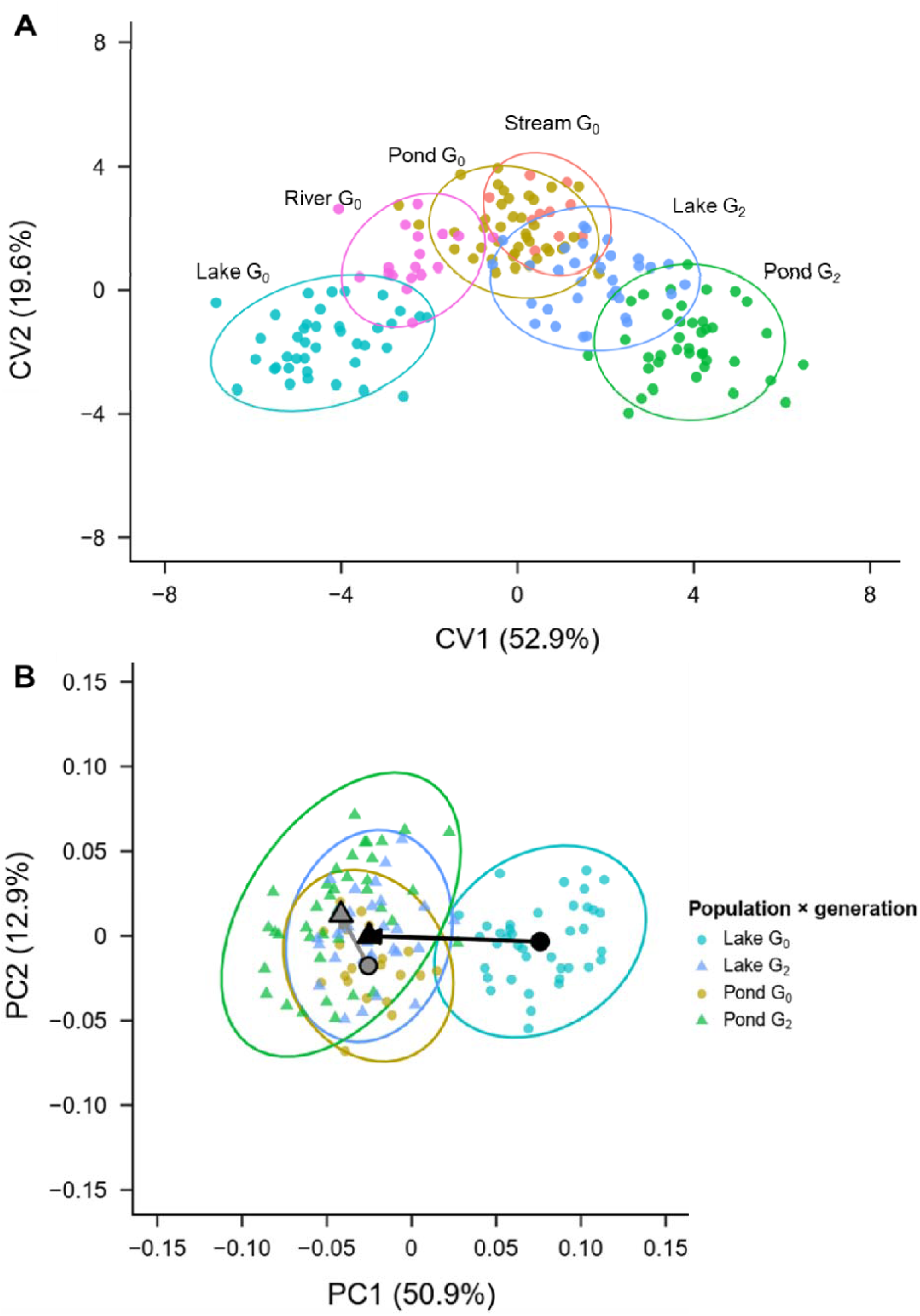
Morphological trajectories across generations in *Pectinidens diaphanus*. (A) Canonical variate analysis of combined G_0_ (wild) and G_2_ (laboratory-reared) individuals from lake and pond populations. Scatterplot of individual scores along CV1 and CV2 derived from Procrustes-aligned landmark coordinates. Points are colored by population of origin and grouped by generation. Ellipses enclose approximately 95% of the individuals of each group. (B) Phenotypic trajectories between wild (G_0_, circles) and laboratory-reared (G_2_, triangles) lake and pond populations in the space of the first two principal components (PC1 and PC2) of shell shape (N = 131). Large symbols are group means and arrows connect the G_0_ to the G_2_ mean of each population. Ellipses enclose 95% of the individuals expected under a bivariate normal distribution within each group.

Shell shape differed significantly between population origins and generations (Procrustes ANOVA; population origin: R^2^ = 0.181, *F*_1, 126_ = 45.34, *P* < 0.001; generation: R^2^ = 0.180, *F*_1, 126_ = 45.18, *P* < 0.001; Table S7A). Centroid size explained 4.2% of shape variation (*F*_1, 126_ = 10.43, *P* < 0.001). The significant interaction indicated that shape differences between G_0_ and G_2_ varied in magnitude between the two population origins (R^2^ = 0.094, *F*_1, 126_ = 23.58, *P* < 0.001). Shell shape differed significantly between G_0_ and G_2_ in both lineages (pairwise comparisons, *P*_Holm_ < 0.001 for both; Table S7B), with a larger effect in the lake lineage. The magnitude of differentiation between lake- and pond-derived snails was lower in G_2_ than in G_0_(pairwise R² = 0.104 vs. 0.496; Table S7B).

Shell shape changed more between G_0_ and G_2_ in the lake-derived population (path length = 0.108) than in the pond-derived population (path length = 0.048; PTA, *d* = 0.060, *Z* = 4.32, *P* < 0.001; Table S7C). The direction of shape changed also differed significantly between the two populations (PTA, vector correlation *r* = 0.426, angle = 64.9°, *Z* = 5.99, *P* < 0.001; Table S7C), indicating that shell shape changed along different multivariate trajectories in the lake-derived and pond-derived populations (Fig. 5B). Lake-derived snails shifted primarily along PC1, whereas pond-derived snails shifted primarily along PC2. Despite these different trajectories, both lineages shifted toward more similar shell morphologies under laboratory conditions (Fig. 5A, B).

## Discussion

This study expands the known range of *P. diaphanus*, the only lymnaeid detected across the surveyed waterbodies in southern Patagonia, which encompass contrasting water flow and permanence. Shell shape differed among wild populations, whereas reproductive anatomy remained conserved. Differences in shell shape between lake- and pond-derived lineages persisted after two generations under common-garden conditions, although they were reduced as both lineages shifted toward more similar laboratory morphotypes. Together, these findings indicate that shell differentiation among populations persists beyond immediate environmental effects, consistent with a heritable component of phenotypic variation.

### A broader distribution and its epidemiological consequences

Our field surveys across 190 sites revealed that *P. diaphanus* has a broader distribution than previously documented. Confirmed records now extend from Santa Cruz Province to Tierra del Fuego, with occurrences at 37 sites compared with the three localities previously reported (Duffy et al. 2009; Correa et al. 2010; Bargues et al. 2012). This discrepancy likely reflects limited previous sampling in a geographically remote and ecologically diverse region rather than a recent range expansion. *P. diaphanus* was not detected alongside other lymnaeid species at the surveyed sites in southern Patagonia.

The distributions of *P. diaphanus* and *G. viator* showed latitudinal segregation, with a narrow zone of potential overlap near the boundary between their ranges. In Argentina, *G. viator* appears to be restricted to northern and central Patagonia and the Pampas region, with no confirmed occurrences south of approximately 42°S (Alda et al. 2021), whereas *P*. *diaphanus* was detected from that latitude southward to Tierra del Fuego. We also recorded *Pseudosuccinea columella* at a single locality in Río Negro Province, extending its known southern distribution in Argentina beyond its previously reported occurrence in Sierra de la Ventana, Buenos Aires Province (Martín et al. 2016). Whether the latitudinal segregation between *P. diaphanus* and *G. viator* reflects environmental differences, biotic interactions, or historical dispersal limitation remains unclear. This distributional pattern contrasts with that of chilinid snails, which occur throughout the Andean region and overlap geographically with the lymnaeid species recorded in our surveys. Targeted sampling across the transition zone, combined with ecological niche modeling and population genetic analyses, could help evaluate the relative roles of environmental conditions, dispersal history, and biotic interactions in shaping these distributions.

The observed distribution has epidemiological implications because *P. diaphanus* was the only lymnaeid species detected at the surveyed sites and therefore represents the sole candidate intermediate host of *F*. *hepatica* in these systems. Defining its geographic range helps identify where this potential intermediate host occurs. If *P*. *diaphanus* is confirmed as a competent intermediate host, its broad occurrence could influence the potential distribution of fasciolosis in southern Patagonia.

### Conserved reproductive anatomy and shell shape differentiation among populations

*Pectinidens diaphanus* occupies both lentic (lakes and ponds) and lotic (rivers and streams) systems, which differ in flow and water permanence (Wetzel 2001). Despite this environmental breadth and the shell variation among populations, reproductive anatomical structures were conserved across populations. This finding supports the taxonomic value of reproductive anatomy in lymnaeids. Although shell morphology can vary substantially (Bonel et al. 2023), copulatory traits tend to be more evolutionarily conserved across environments in Lymnaeids (Pointier et al. 2007; Vinarski and Pointier 2023). The conservation of these structures may reflect functional constraints associated with reproduction (Arnqvist 1998; Hosken and Stockley 2004), although the mechanisms underlying this pattern were not tested in our study. In contrast, variation in shell shape among populations has widely been reported in freshwater gastropods (Dillon 2000; Bourdeau et al. 2015; Whelan 2021). For instance, in *Ampullaceana balthica*, individuals from lotic waterbodies had larger apertures relative to shell length than those from lentic environments, a difference hypothesized to enhance attachment under flowing conditions (Lam and Calow 1988). In our study, the lake population (permanent lentic) had more globose shells and larger apertures than the river population (permanent lotic).

The morphological differences among populations did not align simply with the lotic/lentic distinction but appeared to be associated with differences in water permanence. Ponds and streams in southern Patagonia are subject to recurrent drying (Mayr et al. 2007; Fovet et al. 2021), which may favor morphological traits that reduce water loss during emersion. A smaller aperture reduces the area exposed to evaporation, while narrower shell profiles may facilitate partial burrowing into soft substrates, further reducing exposure to desiccation (Facon et al. 2004; Poznańska et al. 2015). Experimental evidence from other lymnaeids also indicates enhanced desiccation resistance in populations from temporary waterbodies (Chapuis and Ferdy 2012). In the populations examined here, pond-derived snails had more elongated shells and smaller apertures than those from the lake population, traits potentially associated with reduced desiccation risk.

Variation in life-history strategies associated with water permanence may further contribute to these morphological differences. In temporary waterbodies, population persistence depends on the ability to reproduce before desiccation and to recover rapidly after water returns. These conditions may favor a fast-living strategy, characterized by rapid growth, early maturation, and high reproductive investment, consistent with the classical concept of *r*-selected traits (Cole 1954; Stearns 1976; Roff 1993). Differences in growth rate and developmental trajectories can alter shell geometry: faster vertical accretion relative to lateral expansion may contribute to narrower, high-spired morphologies (Chapman 1995; Urdy et al. 2010). Conversely, in permanently inundated environments such as lakes, slower growth trajectories and extended developmental periods may allow greater lateral expansion, resulting in more globose shell allometries (Arthur 1982; Lam and Calow 1988). These explanations are therefore not mutually exclusive: elongated shells with smaller apertures may reflect accelerated growth dynamics associated with temporary waterbodies and confer advantages under desiccation conditions.

The sampled populations span contrasting flows and water permanence, allowing us to examine whether shell morphology covaries with these environmental differences. Shell morphology in freshwater gastropods can vary in response to environmental conditions and biotic interactions, including desiccation risk, predation, and parasite load (Bourdeau et al. 2015; Tariel et al. 2020; Bonel et al. 2021). The observed patterns were consistent with a possible contribution of water permanence and desiccation risk to shell differentiation, although life-history variation and other factors may also contribute. Our sampling design included only one population from each of three waterbody types (lake, river, and stream) and two populations from temporary ponds. This limited replication prevents us from disentangling variation associated with environmental conditions from population-specific differences.

### Persistence of shell differentiation under common-garden conditions

Shell shape differences between lake- and pond-derived lineages persisted after two generations of common-garden rearing, consistent with a heritable component of population-level differentiation in *P*. *diaphanus*. However, the magnitude of this differentiation was reduced in G_2_ compared with G_0_, as both lineages shifted toward more similar, though not identical, laboratory morphotypes. The magnitude of change was greater in the lake-derived lineage, and the two lineages followed significantly different trajectories in morphospace. Thus, the changes associated with common-garden rearing were population-dependent and involved different directions of shape variation. This pattern suggests that environmentally induced variation, potentially involving phenotypic plasticity, may account for a large part of the shell-shape differences observed in wild populations, and can override smaller, genetically based differences. More broadly, the axis of plastic variation may differ from that of genetic differentiation, consistent with the view that plasticity can either shield genotypes from selection or expose them to it (Ghalambor et al. 2007). Comparable patterns have been reported in other gastropods. In *A. balthica*, populations from lotic waterbodies shifted toward the lentic phenotype in the laboratory, whereas lentic populations retained their shell shape (Lam and Calow 1988). In the marine snail *Littorina saxatilis*, ecotype-associated shell traits persisted under common-garden conditions and were associated with genetic differentiation despite ongoing gene flow (Conde-Padín et al. 2009). Whether this larger shift also reflects a stronger plastic response remains unclear.

Several non-exclusive factors could favor greater plasticity in the lake-derived lineage. Conditions likely fluctuate in both lakes and ponds, albeit around different means, which may favor the retention of plasticity in both (Via and Lande 1985). In lakes, plasticity could be further favored if individuals experience microhabitats ranging from areas almost unexposed to desiccation to margins as exposed as ponds, allowing snails to perform well under both conditions (Sultan and Spencer 2002). However, this would require substantial movement or gene flow among these zones, which we did not assess, and all lake individuals were sampled at the margins. Moreover, highly plastic traits may not need to evolve fixed genetic differences, which could also explain why the residual differences in G_2_ are small relative to those in the wild. By contrast, the persistence of differences between lineages may reflect genetic differentiation: the sparse nature of Patagonian waterbodies limits gene flow, and lakes and ponds may be completely disconnected from rivers, allowing populations to follow independent evolutionary trajectories through genetic drift or local selection, although we did not measure gene flow. Testing these hypotheses will require replicated populations and experiments exposing individuals to contrasting environmental conditions. Because offspring were not reared and analyzed as separate families of known parentage, our design cannot separate additive genetic variance from other intergenerational effects. A quantitative genetic study using an appropriate breeding design and pedigree information will be needed to characterize the genetic architecture of shell variation in this species.

## Conclusion

Our study expands the known range of *P. diaphanus* in southern Patagonia and reveals shell differentiation among populations inhabiting environmentally contrasting waterbodies. Shell shape differences among wild populations were consistent with a contribution of water permanence and desiccation risk. Differences between lake- and pond-derived populations persisted under common-garden conditions, although they were reduced as both lineages shifted toward more similar laboratory morphotypes. This pattern suggests that phenotypic plasticity contributes to shell differentiation in wild populations, while differences between experimental populations are consistent with a heritable component of this variation. Together, these findings highlight the importance of distinguishing environmentally induced variation from persistent population differences to understand how intraspecific diversity may contribute to responses to environmental change. Future studies are needed to disentangle the contributions of genetic differentiation and phenotypic plasticity to this variation and to assess their implications for population persistence under increasingly variable climatic conditions.

## Authors’ contributions

Conceptualization, NB and PA; developing methods, NB, MMB, PA, EC, NJ, DF, PJ, PD, JPP; conducting the research, NB, MMB, MA, and PA; data analysis, NB, MMB, PA, MA; data interpretation, NB, MMB, MA, PA; preparation figures & tables, NB, MMB, PA, MA; writing, NB, MMB, PA, EC, SHB, PJ, PD. All authors have read and agreed to the published version of the manuscript.

## Acknowledgments

We would like to thank Antonio Vázquez for his valuable advice and insightful preliminary discussions. We are grateful to the members of Matías Abafatori’s bachelor’s committee (Nicolás Tamburi and Emilia Seuffert) and Micaela Müller’s PhD committee (Benjamin Gourbal, Mathilde Dufay, Annia Alba-Ménendez, Thierry Lefèvre, and Emmanuel Douzery) for their valuable comments and constructive discussions throughout the project. We also thank Rodrigo Gallardo and Amanda Manero for their assistance with snail sampling in Santa Cruz Province. AI-assisted tools were used to improve language and readability and to assist with identifying and correcting errors in *R* scripts. All scientific interpretations and decisions were made by the authors, who take full responsibility for the manuscript.

## Data Availability Statement

The data that support the findings of this study will be made available in the Dryad Digital Repository upon publication of this paper: https://datadryad.org/stash.

