## Supplementary information for "Heritable shell differentiation among populations of the sole lymnaeid snail across freshwaters in southern Patagonia"

#### Table of Contents

***Supplementary Tables***

*Table S1.* Field survey records across freshwater sites in Southern Patagonia 1
*Table S2*. Populations of *Pectinidens diaphanus* selected for morphometric analysis 8
*Table S3.* Morphometric estimates of penial complex and prostate 9
*Table S4.* Results of linear models for penial complex and prostate morphometry 10
*Table S5.* Results of Procrustes ANOVA and pairwise comparisons – $G_0$ 11
*Table S6.* Results of Procrustes ANOVA – $G_2$ 12
*Table S7.* Results of linear models for penial complex and prostate morphometry 13

***Supplementary Figures***

*Figure S1*. Sampled waterbody types 14
*Figure S2*. Experimental breeding design 15
*Figure S3*. Reproductive system anatomy of an adult *Pectinidens diaphanus* 17
*Figure S4.* TCS haplotype network 18

### Supporting tables

**Table S1.** Summary of field surveys conducted at 190 freshwater sites across southern Argentina and Chile during seven field campaigns carried out between 2004 and 2026 in Patagonia. Several sites were revisited during successive sampling campaigns. At each sampling site, GPS coordinates were recorded (UTM X and UTM Y), and the presence or absence of gastropods were determined. When present, the genus of gastropods observed is indicated. Lymnaeids were distinguished from other regional freshwater snails based on shell morphology. A dash (-) indicates that no gastropods were observed at the site. For *Pectinidens diaphanus*, the number of individuals collected (N) was recorded.

| Country | Locality | Habitat | UTMX | UTMY | Date | Gastropod found (Genera) | N |
|---|---|---|---|---|---|---|---|
| Argentina | 7km au sud de Viedma (entrée INTA) | stream | -40.805 | -63.078 | 21/04/2004 | Heleobia | |
| Argentina | Arroyo Blanco | stream | -42.219 | -71.413 | 01/03/2022 | Galba, Chilina | |
| Argentina | General Frias (29km de General Conesa) | stream | -40.249 | -63.181 | 22/04/2004 | Galba, Biomphalaria | |
| Argentina | Lago Escondido Tierra del Fuego | lake | -54.638 | -67.766 | 06/12/2005 | Pectinidens | 9 |
| Argentina | Laguna Coipo | lake | -36.584 | -69.832 | 03/12/2017 | Galba | |
| Argentina | Mechanquil | river | -36.219 | -69.670 | 03/12/2017 | - | |
| Argentina | Balneario Río Limay (Plottié) | river | -38.966 | -68.214 | 04/12/2017 | - | |
| Argentina | Canal Cenillosa | stream | -38.991 | -68.388 | 04/12/2017 | - | |
| Argentina | Canal Riego | stream | -38.988 | -68.390 | 04/12/2017 | - | |
| Argentina | Las Lajas | stream | -38.520 | -70.361 | 04/12/2017 | Galba, Physa, Biomphalaria | |
| Argentina | Manzano Grande | stream | -38.790 | -70.226 | 04/12/2017 | Galba | |
| Argentina | Río Limay | river | -39.022 | -68.463 | 04/12/2017 | Biomphalaria | |
| Argentina | Canal Busto | stream | -38.999 | -64.154 | 05/12/2017 | - | |
| Argentina | Chichinales | river | -39.146 | -66.930 | 05/12/2017 | Pseudosuccinea | |
| Argentina | La Costa | river | -39.092 | -67.208 | 05/12/2017 | - | |
| Argentina | Laguna de Chimbay | lake | -39.173 | -66.139 | 05/12/2017 | Physa | |
| Argentina | Río Colorado | river | -38.977 | -64.106 | 05/12/2017 | - | |
| Argentina | Río Negro (Allen) | river | -39.032 | -67.843 | 05/12/2017 | Physa | |
| Argentina | Río Negro (Jordan) | river | -38.993 | -67.989 | 05/12/2017 | Physa | |
| Argentina | Villa Regina | river | -39.092 | -67.088 | 05/12/2017 | - | |
| Argentina | Arroyo Lepa | stream | -42.615 | -71.076 | 17/11/2018 | - | |
| Argentina | Arroyo Gama | stream | -53.433 | -68.150 | 22/11/2018 | Pectinidens, Chilina, Heleobia | 10 |
| Argentina | Cholila | stream | -42.442 | -71.417 | 17/11/2018 | Galba | |
| Argentina | Cuesta del arbolito | stream | -43.166 | -70.839 | 17/11/2018 | Galba | |

**Table S1.** *Continued…*

| Country | Locality | Habitat | UTMX | UTMY | Date | Gastropod found (Genera) | N |
|---|---|---|---|---|---|---|---|
| Argentina | El Bolsón | water tank | -41.865 | -71.535 | 17/11/2018 | - | |
| Argentina | La Paulina | river | -43.805 | -70.935 | 17/11/2018 | Galba | |
| Argentina | Leleque | stream | -42.432 | -71.103 | 17/11/2018 | - | |
| Argentina | Arroyo Apeleg | stream | -44.830 | -70.127 | 18/11/2018 | Galba | |
| Argentina | Arroyo Feo | stream | -46.876 | -70.738 | 18/11/2018 | - | |
| Argentina | Arroyo Genoa (La Laurita) | stream | -44.830 | -70.127 | 18/11/2018 | Galba, Biomphalaria | |
| Argentina | Arroyo Page | stream | -46.734 | -70.840 | 18/11/2018 | - | |
| Argentina | Arroyo Ingreso INTA | stream | -51.857 | -70.426 | 01/03/2025 | Chilina | |
| Argentina | Facundo | stream | -45.318 | -69.945 | 18/11/2018 | Galba, Biomphalaria | |
| Argentina | Río Ecker | river | -47.123 | -70.889 | 18/11/2018 | Pectinidens | 30 |
| Argentina | Río fenix (dans la localité de Perito moreno) | river | -46.583 | -70.917 | 18/11/2018 | Galba | |
| Argentina | Río Mayo (Cañada Raquicu) | lake | -45.318 | -69.945 | 18/11/2018 | - | |
| Argentina | Río Pinturas | river | -47.007 | -70.832 | 18/11/2018 | - | |
| Argentina | Antes de la Estancia La Primitiva | seco | -48.732 | -70.525 | 19/11/2018 | - | |
| Argentina | Arroyo Nirihuau | stream | -41.087 | -71.183 | 27/02/2022 | Galba, Chilina | |
| Argentina | El Javi | stream | -48.339 | -70.852 | 19/11/2018 | - | |
| Argentina | Estancia la Lucía | inflow | -49.229 | -71.353 | 19/11/2018 | - | |
| Argentina | Estancia La Primitiva (valle de río chico) | stream | -48.739 | -70.438 | 19/11/2018 | Galba | |
| Argentina | Hacienda La Lucha | flooded meadow | -48.507 | -70.434 | 19/11/2018 | - | |
| Argentina | Puesto Chacra | flooded meadow | -49.558 | -72.415 | 19/11/2018 | - | |
| Argentina | Río Cangrejo | river | -49.591 | -72.266 | 19/11/2018 | - | |
| Argentina | Río Chalia | river | -49.584 | -71.439 | 19/11/2018 | - | |
| Argentina | Río Chico (Estancia Las Horquetas. 50 metros de ancho) | river | -48.269 | -71.166 | 19/11/2018 | - | |
| Argentina | Río Olnie | river | -47.641 | -70.808 | 19/11/2018 | - | |
| Argentina | Arroyo Josefina | stream | -50.319 | -71.956 | 20/11/2018 | - | |
| Argentina | Arroyo Turbio | stream | -50.003 | -72.110 | 20/11/2018 | - | |
| Argentina | El Calafate (parque recreacional) | flooded meadow | -50.339 | -71.289 | 20/11/2018 | - | |
| Argentina | Estancia Buenos Aires | stream | -50.353 | -72.730 | 20/11/2018 | - | |
| Argentina | Lago Argentino a paso Viggieri | lake | -50.124 | -72.085 | 20/11/2018 | - | |
| Argentina | Arroyo Suspiro | stream | -54.274 | -67.205 | 23/11/2018 | Pectinidens, Chilina | 8 |

**Table S1.** *Continued…*

| Country | Locality | Habitat | UTMX | UTMY | Date | Gastropod found (Genera) | N |
|---|---|---|---|---|---|---|---|
| Argentina | Río de las Vueltas | river | -49.311 | -72.899 | 20/11/2018 | Pectinidens | 30 |
| Argentina | Arroyo Telken | river | -46.883 | -70.734 | 24/02/2025 | Chilina | |
| Argentina | Río Mitre | river | -50.400 | -72.733 | 20/11/2018 | - | |
| Argentina | Arroyo cerca de Cerro Sombrero | stream | -51.302 | -72.181 | 21/11/2018 | Pectinidens | 2 |
| Argentina | Arroyo Italiano (coordenadas a 200 metros) | stream | -50.769 | -71.430 | 21/11/2018 | Pectinidens | 2 |
| Argentina | Chorrillo de Barrancas Blancas | stream | -50.875 | -71.490 | 21/11/2018 | Pectinidens | 2 |
| Argentina | Estancia Puentes del Coig | river | -51.143 | -71.951 | 21/11/2018 | - | |
| Argentina | Laguna Condor | lake | -51.758 | -71.760 | 21/11/2018 | - | |
| Argentina | Río Bella Vista | river | -51.865 | -70.559 | 21/11/2018 | Pectinidens | 2 |
| Argentina | Río Bote | river | -50.295 | -71.718 | 21/11/2018 | - | |
| Argentina | Río Coig (Estancia Tapi Aike) | river | -51.019 | -71.777 | 21/11/2018 | - | |
| Argentina | Arroyo Valcheta | stream | -40.679 | -66.175 | 24/02/2022 | Chilina, Heleobia | |
| Argentina | Río Gallegos à Puente Blanco | river | -51.894 | -71.596 | 21/11/2018 | Pectinidens | 20 |
| Argentina | Río Gallegos Güer Aike | river | -51.652 | -69.629 | 21/11/2018 | Pectinidens | 2 |
| Argentina | Río Telque | river | -50.774 | -71.428 | 21/11/2018 | - | |
| Argentina | Río Turbio | river | -51.447 | -72.226 | 21/11/2018 | - | |
| Argentina | Río Turbio (Estancia Stag River) | inflow | -51.686 | -72.004 | 21/11/2018 | Pectinidens | 40 |
| Argentina | Río Turbio à Glen Cross | river | -51.822 | -71.699 | 21/11/2018 | - | |
| Argentina | Bajo Caracoles | stream | -47.123 | -70.889 | 19/11/2018 | Heleobia, Chilina | |
| Chile | Bahía Santiago | stream | -52.485 | -69.849 | 22/11/2018 | - | |
| Argentina | Chorrillo La Misión | water trough | -53.669 | -67.802 | 22/11/2018 | - | |
| Argentina | Bolsón El Paralelo | stream | -41.963 | -71.533 | 17/11/2018 | Chilina | |
| Chile | Estancia Marisol | stream | -53.028 | -68.801 | 22/11/2018 | Pectinidens | 30 |
| Argentina | Río Chico (Estancia Violeta) | river | -53.674 | -67.935 | 22/11/2018 | - | |
| Argentina | Camusu Aike | river | -51.015 | -70.798 | 23/03/2022 | Chilina | |
| Argentina | Arroyo Khami | stream | -54.420 | -67.218 | 23/11/2018 | Pectinidens | 20 |
| Argentina | Cañadón Pinturas | river | -47.142 | -70.675 | 25/02/2025 | Chilina | |
| Chile | Cuarto Chirillo | river | -52.733 | -68.996 | 22/11/2018 | Chilina | |
| Argentina | El Pluma | stream | -46.487 | -70.024 | 24/02/2025 | Galba, Chilina | |
| Argentina | Epuyen cressonnière | stream | -42.221 | -71.358 | 02/03/2022 | Chilina | |

**Table S1.** *Continued…*

| Country | Locality | Habitat | UTMX | UTMY | Date | Gastropod found (Genera) | N |
|---|---|---|---|---|---|---|---|
| Argentina | Estancia Coy Aike | river | -51.153 | -69.525 | 25/03/2022 | Chilina | |
| Argentina | Río Olivia | river | -54.725 | -68.001 | 23/11/2018 | - | |
| Argentina | Río Tristen | river | -54.731 | -67.941 | 23/11/2018 | - | |
| Argentina | Laguna Azul 2018 | lake | -52.075 | -69.581 | 24/11/2018 | Pectinidens | 30 |
| Argentina | Cañada La Zaranda | stream | -46.520 | -67.464 | 25/11/2018 | - | |
| Argentina | cañada Minerales | stream | -46.714 | -67.367 | 25/11/2018 | - | |
| Argentina | Estancia Las Luises | lake | -50.619 | -69.360 | 25/11/2018 | - | |
| Argentina | La Cabaña | lake | -47.954 | -67.541 | 25/11/2018 | - | |
| Argentina | Río Chico à Constanza | river | -49.777 | -68.639 | 25/11/2018 | Pectinidens | 2 |
| Argentina | Río Coyle | river | -51.153 | -69.525 | 25/11/2018 | Pectinidens | 2 |
| Argentina | Río Deseado | river | -47.184 | -67.248 | 25/11/2018 | - | |
| Argentina | Estancia Juanita | stream | -46.084 | -70.518 | 18/11/2018 | Chilina | |
| Argentina | Tehuelches | stream | -46.895 | -67.283 | 25/11/2018 | - | |
| Argentina | Arroyo Genoa à Magnífica rotonda | stream | -44.057 | -70.530 | 26/11/2018 | Pectinidens | 50 |
| Argentina | Estancia La Constancia | river | -49.777 | -68.638 | 25/03/2022 | Chilina | |
| Argentina | Estancia La Jorgelina | stream | -48.774 | -70.424 | 19/03/2022 | Chilina | |
| Argentina | Estancia La Lucia | stream | -49.231 | -71.347 | 19/03/2022 | Chilina | |
| Argentina | Sarmiento Este | flooded meadow | -45.599 | -69.022 | 26/11/2018 | - | |
| Argentina | General Conesa Lago | lake | -40.092 | -64.483 | 23/02/2022 | - | |
| Argentina | Laguna Cortez Tanque Australiano + Bebedero | water trough | -40.522 | -64.691 | 23/02/2022 | - | |
| Argentina | Estancia Menendez (Efluente del Río candelaria) | inflow | -53.857 | -67.768 | 23/11/2018 | Chilina | |
| Argentina | Guardia Mitre (76 km au sud de Viedma - route à Conesa) | stream | -40.436 | -63.679 | 22/04/2004 | Chilina, Heleobia, Anodonta | |
| Argentina | Hacienda Roxana source + mare | stream + pond | -41.030 | -70.238 | 25/02/2022 | Galba | |
| Argentina | Los Menucos pozos y lagunas | pond | -40.850 | -68.182 | 25/02/2022 | - | |
| Argentina | Moutons Tanque australiano +bebederos | taker tank + trough | -41.138 | -68.572 | 25/02/2022 | - | |
| Argentina | Río Pilcaniyeu | river | -41.089 | -70.830 | 25/02/2022 | Galba, Biomphalaria | |
| Argentina | Arroyo Media Luna | stream | -40.629 | -71.486 | 27/02/2022 | - | |
| Argentina | Isla Pavón 2022 | river | -50.000 | -68.930 | 25/03/2022 | Pectinidens, Chilina | 20 |
| Argentina | La Angostura | stream | -40.763 | -71.639 | 27/02/2022 | - | |
| Argentina | José de San martón (Arroyo Genoa) | stream | -44.059 | -70.482 | 26/11/2018 | Chilina | |

**Table S1.** *Continued…*

| Country | Locality | Habitat | UTMX | UTMY | Date | Gastropod found (Genera) | N |
|---|---|---|---|---|---|---|---|
| Argentina | Río Huemel | river | -40.896 | -71.472 | 27/02/2022 | - | |
| Argentina | La Negrita canal | stream | -43.287 | -65.578 | 04/03/2022 | Chilina | |
| Argentina | Río Minero | river | -40.691 | -71.292 | 27/02/2022 | - | |
| Argentina | Bosque del Pino | river | -41.252 | -71.190 | 28/02/2022 | Aroapyrgus | |
| Argentina | El Bolson bassin dans la ville | pond | -41.965 | -71.535 | 28/02/2022 | Physa | |
| Argentina | La Buitrera petit ruisseau | stream | -41.284 | -71.178 | 28/02/2022 | Galba, Aroapyrgus | |
| Argentina | Lago Fagnano | lake | -54.551 | -67.224 | 23/11/2018 | Chilina | |
| Argentina | Río Villegas | river | -41.586 | -71.492 | 28/02/2022 | - | |
| Argentina | Lago Puelo | lake | -42.061 | -71.596 | 02/03/2022 | Chilina | |
| Argentina | Laguna Azul 2026 | lake | -52.077 | -69.585 | 09/03/2026 | Pectinidens, Chilina | 50 |
| Argentina | Las Horquetas 2026 | river | -51.411 | -70.193 | 07/03/2026 | Pectinidens, Chilina | 40 |
| Argentina | Mirador Lago Viedma | lake | -49.717 | -71.96 | 20/11/2018 | Chilina | |
| Argentina | Río Piedrogoso | river | -42.140 | -71.41 | 01/03/2022 | - | |
| Argentina | Confluencia | river | -42.000 | -71.587 | 02/03/2022 | - | |
| Argentina | Playa Bonita Bariloche source | stream | -41.121 | -71.398 | 28/02/2022 | Chilina, Physa | |
| Argentina | Rincon Chico Río | river | -40.991 | -71.104 | 27/02/2022 | Galba, Chilina | |
| Argentina | Río Azul | river | -42.021 | -71.59 | 02/03/2022 | Chilina | |
| Argentina | Río Bajo Grande | river | -42.208 | -71.389 | 01/03/2022 | Galba, Chilina | |
| Argentina | Río Chico | river | -48.752 | -70.323 | 18/03/2022 | Pectinidens, Chilina | 10 |
| Argentina | Arroyo Leleque | stream | -42.537 | -71.101 | 03/03/2022 | - | |
| Argentina | Arroyo Lepa | stream | -42.614 | -71.077 | 03/03/2022 | Galba | |
| Argentina | Cuesta del Arbolio canal | stream | -43.166 | -70.839 | 03/03/2022 | Galba, Biomphalaria | |
| Argentina | INTA Trevelin prairie inondée | flooded meadow | -43.121 | -71.54 | 03/03/2022 | Galba | |
| Argentina | Leleque affluent | inflow | -42.516 | -71.095 | 03/03/2022 | - | |
| Argentina | Río el Zurdo | river | -51.659 | -71.232 | 21/11/2018 | Pectinidens, Chilina | 6 |
| Argentina | Río Epuyen | river | -42.139 | -71.41 | 01/03/2022 | Chilina | |
| Argentina | Río Chubut 1 | river | -43.855 | -68.814 | 04/03/2022 | - | |
| Argentina | Río Chubut 2 | river | -43.833 | -67.902 | 04/03/2022 | - | |
| Argentina | Río Chubut 3 | river | -43.273 | -65.251 | 04/03/2022 | - | |
| Argentina | Río Chubut à Playa Union | river | -43.306 | -65.091 | 04/03/2022 | - | |

**Table S1.** *Continued…*

| Country | Locality | Habitat | UTMX | UTMY | Date | Gastropod found (Genera) | N |
|---|---|---|---|---|---|---|---|
| Argentina | Zanjon Madre affluent | inflow | -43.741 | -67.458 | 04/03/2022 | - | |
| Argentina | Trelew canal | stream | -43.236 | -65.293 | 05/03/2022 | Physa | |
| Argentina | Frias 2022 | stream | -40.260 | -64.190 | 07/03/2022 | Galba | |
| Argentina | La Luisa zone inondée | pond | -40.170 | -64.305 | 07/03/2022 | - | |
| Argentina | Sierra Grande arroyo | stream | -41.598 | -65.354 | 07/03/2022 | - | |
| Argentina | Centro de Investigacion Zoonosis | flooded meadow | -45.592 | -69.023 | 17/03/2022 | - | |
| Argentina | Chacra CEFAM | pond | -45.878 | -69.079 | 17/03/2022 | Galba, Biomphalaria | |
| Argentina | Chacra Oscar Jensen | pond | -45.592 | -69.043 | 17/03/2022 | Galba | |
| Argentina | Machin Casa Laura | pond | -45.592 | -69.037 | 17/03/2022 | - | |
| Argentina | Cañadon cantado | stream | -45.389 | -69.99 | 18/03/2022 | - | |
| Argentina | Estancia Santa Thelma | stream | -45.726 | -70.272 | 18/03/2022 | - | |
| Argentina | Río Epuyen Las Pitras | river | -42.084 | -71.527 | 02/03/2022 | Chilina | |
| Argentina | Río Ewan Norte | river | -54.125 | -67.276 | 23/11/2018 | Pectinidens, Chilina | 30 |
| Argentina | Río Fuego | river | -53.978 | -67.456 | 23/11/2018 | Chilina | |
| Argentina | Gobernador Gregores | river | -48.753 | -70.261 | 19/03/2022 | Pectinidens | 30 |
| Argentina | Río La Leona (hotel La Leona) | river | -49.808 | -72.053 | 20/11/2018 | Chilina | |
| Argentina | El Chalten | stream | -49.332 | -72.92 | 20/03/2022 | - | |
| Argentina | Laguna Torre | lake | -49.330 | -72.990 | 20/03/2022 | - | |
| Argentina | Avant Estancia Las Horquetas | stream | -51.399 | -70.217 | 23/03/2022 | Potamolythus | |
| Argentina | Río Limay | river | -41.056 | -71.148 | 27/02/2022 | Chilina | |
| Argentina | Cañadon El Calafate | river | -50.355 | -72.242 | 23/03/2022 | Pectinidens | 30 |
| Argentina | Las Horquetas 2022 | river | -51.412 | -70.194 | 23/03/2022 | Pectinidens | 20 |
| Argentina | Río Bote | river | -50.295 | -71.718 | 23/03/2022 | Pectinidens | 3 |
| Argentina | Chorrillo de los Frailes 2022 | pond | -51.907 | -69.127 | 24/03/2022 | Pectinidens | 34 |
| Argentina | Estancia El Condor | stream | -51.927 | -69.12 | 24/03/2022 | Cochliopidae | |
| Argentina | Laguna Azul 2022 | lake | -52.076 | -69.58 | 24/03/2022 | Pectinidens | 17 |
| Argentina | Río Minas | river | -42.216 | -71.356 | 01/03/2022 | Galba, Chilina | |
| Argentina | Río Quemquemtreu | river | -42.000 | -71.570 | 02/03/2022 | Chilina | |
| Argentina | Río Santa cruz (Isla Pavón) | river | -50.002 | -68.928 | 25/11/2018 | Pectinidens, Chilina | 30 |
| Argentina | Río seco | dry ditch | -48.521 | -67.730 | 26/03/2022 | - | |

**Table S1.** *Continued…*

| Country | Locality | Habitat | UTMX | UTMY | Date | Gastropod found (Genera) | N |
|---|---|---|---|---|---|---|---|
| Argentina | Cancha Carrera | stream | -51.142 | -71.950 | 09/11/2023 | Pectinidens | 20 |
| Argentina | Corpen Aike | pond | -51.386 | -70.237 | 09/11/2023 | Pectinidens | 40 |
| Argentina | Fuentes del Coig | stream | -51.142 | -71.950 | 09/11/2023 | Pectinidens | 15 |
| Argentina | la Vanguardia | flooded meadow | -51.031 | -71.045 | 09/11/2023 | - | |
| Argentina | Las Horquetas 2023 | river | -51.412 | -70.193 | 09/11/2023 | Pectinidens | 25 |
| Argentina | Tapi Aike (puesto fijo) | river | -51.058 | -71.803 | 09/11/2023 | Pectinidens | 10 |
| Argentina | Laguna Potrok Aike 2023 | lake | -51.955 | -70.397 | 12/11/2023 | Pectinidens | 20 |
| Argentina | Laguna Potrok Aike source 2023 | stream | -51.968 | -70.422 | 12/11/2023 | Pectinidens | 15 |
| Argentina | Río turbio à estancia Glencross | river | -51.822 | -71.699 | 12/11/2023 | - | |
| Argentina | Santa Ana | river | -51.686 | -71.975 | 12/11/2023 | Pectinidens | 5 |
| Argentina | Chimen Aike | stream | -51.709 | -69.287 | 13/11/2023 | - | |
| Argentina | Laguna Azul 2023 | lake | -52.077 | -69.585 | 13/11/2023 | Pectinidens | 20 |
| Argentina | Chorrillo de los Frailes 2023 | pond | -51.906 | -69.127 | 14/11/2023 | Pectinidens | 30 |
| Argentina | Río Senguer (à Colhuehuapi) | river | -45.676 | -68.914 | 26/11/2018 | Chilina | |
| Argentina | Río Senguer (à Sarmiento) | river | -45.765 | -69.183 | 26/11/2018 | Chilina | |
| Argentina | Río Fenix à Perito Moreno | river | -46.583 | -70.917 | 24/02/2025 | Cochliopidae | |
| Chile | Río Side (Estancia Los Mellizos) | river | -52.730 | -69.240 | 22/11/2018 | Pectinidens, Chilina | 1 |
| Argentina | Bajo Caracoles | river | -47.444 | -70.925 | 26/02/2025 | Pectinidens | 20 |
| Argentina | Laguna de Los Tres | lake | -49.279 | -72.985 | 27/02/2025 | - | |
| Argentina | Laguna Madie | lake | -49.298 | -72.953 | 27/02/2025 | - | |
| Argentina | Río Tecka | river | -43.463 | -70.856 | 03/03/2022 | Chilina | |
| Argentina | Laguna Potrok Aike 2025 | lake | -51.955 | -70.397 | 01/03/2025 | Pectinidens | 50 |
| Argentina | Potrok Aike source | stream | -51.968 | -70.405 | 01/03/2025 | - | |
| Argentina | Frias 2025 | stream | -40.249 | -64.182 | 09/05/2025 | Galba, Biomphalaria, Physa | |
| Argentina | Cancha Carrera | stream | -51.143 | -71.951 | 07/03/2026 | Pectinidens | 30 |
| Argentina | Tres Lagos | river | -49.597 | -71.447 | 19/03/2022 | Pectinidens, Chilina | 21 |
| Argentina | Cerro Cuadrado | pond | -51.671 | -71.918 | 08/03/2026 | Pectinidens | 40 |
| Argentina | Laguna Potrok Aike 2026 | lake | -51.956 | -70.396 | 08/03/2026 | Pectinidens | 35 |
| Argentina | Tunquelen canal | stream | -40.492 | -67.256 | 24/02/2022 | Chilina | |
| Argentina | Chorrillo de los Frailes 2026 | pond | -51.907 | -69.111 | 10/03/2026 | Pectinidens | 30 |
| Argentina | Frias 2026 | stream | -40.249 | -64.182 | 13/05/2026 | Galba, Biomphalaria, Physa | |

**Table S2.** Populations of *Pectinidens diaphanus* selected for morphometric analysis in Patagonia. To investigate morphological variation among populations, five representative populations were selected from four contrasting habitat types. All populations were sampled in November 2023 and are located at similar latitudes in southern Santa Cruz Province, Argentina. N: number of individuals photographed.

| Habitat | Site | Latitude | Longitude | N |
|---|---|---|---|---|
| Lake | Laguna Azul | -52.078 | -69.586 | 39 |
| River | Las Horquetas | -51.412 | -70.193 | 18 |
| Stream | Cancha Carrera | -51.142 | -71.950 | 10 |
| Pond | Corpen Aike | -51.387 | -70.237 | 20 |
| | Chorrillo de los Frailes | -51.906 | -69.128 | 20 |

**Table S3.** Morphometry of the reproductive system of wild populations. Mean values (± S.D.) of the area ($mm^2$) and perimeter (mm) of the penial complex and prostate of the lymnaeid snail *Pectinidens diaphanus* collected in Patagonia. Morphometric traits were quantified using *ImageJ* software. All morphometric traits were compared among habitat types using ANOVA in *R Studio*. We found no differences in penial complex or prostate area and perimeter among habitat types (see Table S4 for statistics details). N: number of individuals measured.

| Population origin | Penial complex | | Prostate | | N |
|---|---|---|---|---|---|
| | Area | Perimeter | Area | Perimeter | |
| Lake | 3.05 ± 0.97 | 11.56 ± 1.95 | 3.04 ± 0.79 | 6.72 ± 0.90 | 10 |
| River | 2.29 ± 0.31 | 10.3 ± 0.67 | 2.19 ± 0.41 | 5.57 ± 0.52 | 10 |
| Stream | 3.61 ± 0.63 | 13.21 ± 0.78 | 2.89 ± 0.47 | 6.37 ± 0.45 | 10 |
| Pond | 2.79 ± 0.69 | 11.49 ± 0.99 | 2.5 ± 0.48 | 5.79 ± 0.48 | 20 |

**Table S4.** Statistical significance of the ANOVA testing for population origin effect on area and perimeter of the penial complex and prostate of the lymnaeid snail *Pectinidens diaphanus* collected in Patagonia. Population effect sizes were tested using multiple pairwise comparisons with the Holm-Bonferroni correction for multiple testing when ANOVA results were significant.

| | Traits measured | Population effect | Population comparison | Effect size |
|---|---|---|---|---|
| Prostate | Area | $F_{(3, 46)} = 1.86$ | – | – |
| | Perimeter | $F_{(3, 46)} = 3.30$ * | Lake-River | 2.67 |
| | | | Lake-Pond | 2.49 |
| | | | River-Stream | -1.87 |
| | | | Pond-Stream | -1.57 |
| | | | Lake-Stream | 0.80 |
| | | | River-Pond | -0.59 |
| Penial Complex | Area | $F_{(3, 46)} = 4.04$ * | River-Stream | -3.38 |
| | | | Pond-Stream | -2.42 |
| | | | Lake-River | 1.95 |
| | | | River-Pond | -1.49 |
| | | | Lake-Stream | -1.44 |
| | | | Lake-Pond | 0.76 |
| | Perimeter | $F_{(3, 46)} = 5.35$ ** | River-Stream | -3.97 * |
| | | | Pond-Stream | -2.70 |
| | | | Lake-Stream | -2.24 |
| | | | River-Pond | -1.89 |
| | | | Lake-River | 1.73 |
| | | | Lake-Pond | 0.11 |

* $P < 0.05$, ** $P < 0.01$.

**Table S5.** Overall Procrustes ANOVA (A) and pairwise comparisons (B) testing shell shape differences among wild populations ($G_0$) of *Pectinidens diaphanus* from contrasting freshwaters in southern Patagonia. Shape variation was tested as a function of population origin while controlling for centroid size as an allometric covariate. Effect size is represented by $R^2$, indicating the proportion of shape variation explained by population differences. Significance was assessed using 9,999 residual permutations. *P* values for pairwise comparisons were adjusted using Holm's method.

**A.** Overall Procrustes ANOVA

| **Factor** | **df** | **Sums of Sq** | **Mean Sq** | $R^2$ | ***F*-value** | ***P*-value** |
|---|---|---|---|---|---|---|
| Centroid size (logCS) | 1 | 0.017 | 0.017 | 0.022 | 4.67 | < 0.001 |
| Population origin | 3 | 0.257 | 0.086 | 0.493 | 34.53 | < 0.001 |
| Residuals | 102 | 0.253 | 0.002 | 0.485 | – | – |
| Total | 106 | 0.522 | – | 1 | – | – |

**B.** Pairwise Procrustes ANOVA Comparisons

| **Population comparison** | **df** | ***F*-value** | ***P*-value** | ***P* (Holm-adjusted)** |
|---|---|---|---|---|
| Stream vs. Pond | 1, 47 | 3.51 | 0.0013 | 0.0013 |
| Stream vs. Lake | 1, 46 | 38.47 | < 0.001 | < 0.001 |
| Stream vs. River | 1, 25 | 12.98 | < 0.001 | < 0.001 |
| Pond vs. Lake | 1, 76 | 79.19 | < 0.001 | < 0.001 |
| Pond vs. River | 1, 55 | 14.80 | < 0.001 | < 0.001 |
| Lake vs. River | 1, 54 | 17.47 | < 0.001 | < 0.001 |

**Table S6.** Overall Procrustes ANOVA (A) and pairwise comparisons (B) testing shell-shape differences among six population–generation groups of *Pectinidens diaphanus* from contrasting freshwaters in southern Patagonia. These groups comprised wild ($G_0$) stream, pond, lake and river snails, and laboratory-reared ($G_2$) pond- and lake-derived snails. Shape variation was tested as a function of population origin while controlling for centroid size as an allometric covariate. Effect size is represented by $R^2$, indicating the proportion of shape variation explained by each model term or group contrast. Significance was assessed using 9,999 residual permutations. *P* values for pairwise comparisons were adjusted using Holm's method.

**A.**

| Factor | df | Sums of Sq | Mean Sq | $R^2$ | *F*-value | Z-score | *P*-value |
|---|---|---|---|---|---|---|---|
| Centroid size | 1 | 0.016 | 0.016 | 0.017 | 5.55 | 3.90 | < 0.001 |
| Population origin | 5 | 0.439 | 0.088 | 0.468 | 31.21 | 9.59 | < 0.001 |
| Residuals | 172 | 0.483 | 0.003 | 0.516 | – | – | – |
| Total | 178 | 0.938 | – | 1 | – | – | – |

**B.**

| Population comparison | df | $R^2$ | *F*-value | *P*-value | *P* (Holm-adjusted) |
|---|---|---|---|---|---|
| Stream ($G_0$) vs. Pond ($G_0$) | 1, 47 | 0.068 | 3.52 | 0.0013 | 0.0015 |
| Stream ($G_0$) vs. Pond ($G_2$) | 1, 46 | 0.118 | 7.17 | < 0.001 | 0.0015 |
| Stream ($G_0$) vs. Lake ($G_0$) | 1, 46 | 0.437 | 38.46 | < 0.001 | 0.0015 |
| Stream ($G_0$) vs. Lake ($G_2$) | 1, 40 | 0.087 | 4.10 | < 0.001 | 0.0015 |
| Stream ($G_0$) vs. River ($G_0$) | 1, 25 | 0.326 | 12.99 | < 0.001 | 0.0015 |
| Pond ($G_0$) vs. Pond ($G_2$) | 1, 76 | 0.172 | 17.86 | < 0.001 | 0.0015 |
| Pond ($G_0$) vs. Lake ($G_0$) | 1, 76 | 0.496 | 79.15 | < 0.001 | 0.0015 |
| Pond ($G_0$) vs. Lake ($G_2$) | 1, 70 | 0.144 | 12.34 | < 0.001 | 0.0015 |
| Pond ($G_0$) vs. River ($G_0$) | 1, 55 | 0.205 | 14.81 | < 0.001 | 0.0015 |
| Pond ($G_2$) vs. Lake ($G_0$) | 1, 75 | 0.510 | 82.23 | < 0.001 | 0.0015 |
| Pond ($G_2$) vs. Lake ($G_2$) | 1, 69 | 0.104 | 8.85 | < 0.001 | 0.0015 |
| Pond ($G_2$) vs. River ($G_0$) | 1, 54 | 0.300 | 26.78 | < 0.001 | 0.0015 |
| Lake ($G_0$) vs. Lake ($G_2$) | 1, 69 | 0.471 | 64.50 | < 0.001 | 0.0015 |
| Lake ($G_0$) vs. River ($G_0$) | 1, 54 | 0.230 | 17.45 | < 0.001 | 0.0015 |
| Lake ($G_2$) vs. River ($G_0$) | 1, 48 | 0.233 | 15.38 | < 0.001 | 0.0015 |

**Table S7.** Procrustes ANOVA and pairwise comparisons of shell shape between wild ($G_0$) and laboratory-reared ($G_2$) lake and pond populations of *Pectinidens diaphanus* (N = 131; lake: $G_0$ n = 39, $G_2$ n = 33; pond: $G_0$ n = 20, $G_2$ n = 39). (A) Factorial Procrustes ANOVA with log-transformed centroid size (logCS) as an allometric covariate, population origin (pond vs. lake), generation ($G_0$ vs. $G_2$) and their interaction, using sequential (type I) sums of squares in the order shown. $R^2$ is the proportion of total shape variation explained by each term. (B) Pairwise Procrustes ANOVAs between the four population × generation groups (with shape ~ group + logCS, entering group first). $R^2$ is the proportion of variation explained by group. (C) Phenotypic trajectory analysis (PTA) of the $G_0 \rightarrow G_2$ shape change. Significance was assessed with 10,000 residual permutations; raw and Holm-adjusted P values are reported in (B).

**A.**

| Factor | df | Sums of Sq | Mean Sq | $R^2$ | *F*-value | Z-score | *P*-value |
|---|---|---|---|---|---|---|---|
| Centroid size | 1 | 0.032 | 0.032 | 0.042 | 10.43 | 3.48 | < 0.001 |
| Population origin | 1 | 0.140 | 0.140 | 0.181 | 45.34 | 5.06 | < 0.001 |
| Generation | 1 | 0.140 | 0.140 | 0.180 | 45.18 | 5.72 | < 0.001 |
| Pop × Gen | 1 | 0.073 | 0.073 | 0.094 | 23.58 | 5.78 | < 0.001 |
| Residuals | 126 | 0.389 | 0.003 | 0.503 | – | – | – |
| Total | 130 | 0.773 | – | 1 | – | – | – |

**B.**

| Population comparison | df | $R^2$ | *F*-value | *P*-value | *P* (Holm-adjusted) |
|---|---|---|---|---|---|
| Pond $G_0$ – Pond $G_2$ | 1, 56 | 0.136 | 10.22 | < 0.001 | < 0.001 |
| Pond $G_0$ – Lake $G_0$ | 1, 56 | 0.459 | 50.94 | < 0.001 | < 0.001 |
| Pond $G_0$ – Lake $G_2$ | 1, 50 | 0.115 | 6.94 | < 0.001 | < 0.001 |
| Pond $G_2$ – Lake $G_0$ | 1, 75 | 0.510 | 82.18 | < 0.001 | < 0.001 |
| Pond $G_2$ – Lake $G_2$ | 1, 69 | 0.104 | 8.85 | < 0.001 | < 0.001 |
| Lake $G_0$ – Lake $G_2$ | 1, 69 | 0.471 | 64.48 | < 0.001 | < 0.001 |

**C.**

| Attribute | Lake | Pond | Statistic | Z-score | *P* |
|---|---|---|---|---|---|
| Trajectory length | 0.108 | 0.048 | d = 0.061 | 4.32 | < 0.001 |
| Direction | | | r = 0.426; angle = 64.9° | 5.99 | < 0.001 |

#### Supporting figures

**Figure S1.** Populations sampled for morphometric analysis. (A) Classification of the four habitat types from which *Pectinidens diaphanus* populations were collected: river and stream (permanent and temporary lotic environments), and lake and small pond (permanent and temporary lentic environments). (B) Geographic location of the populations sampled in southern Santa Cruz province, Argentina.

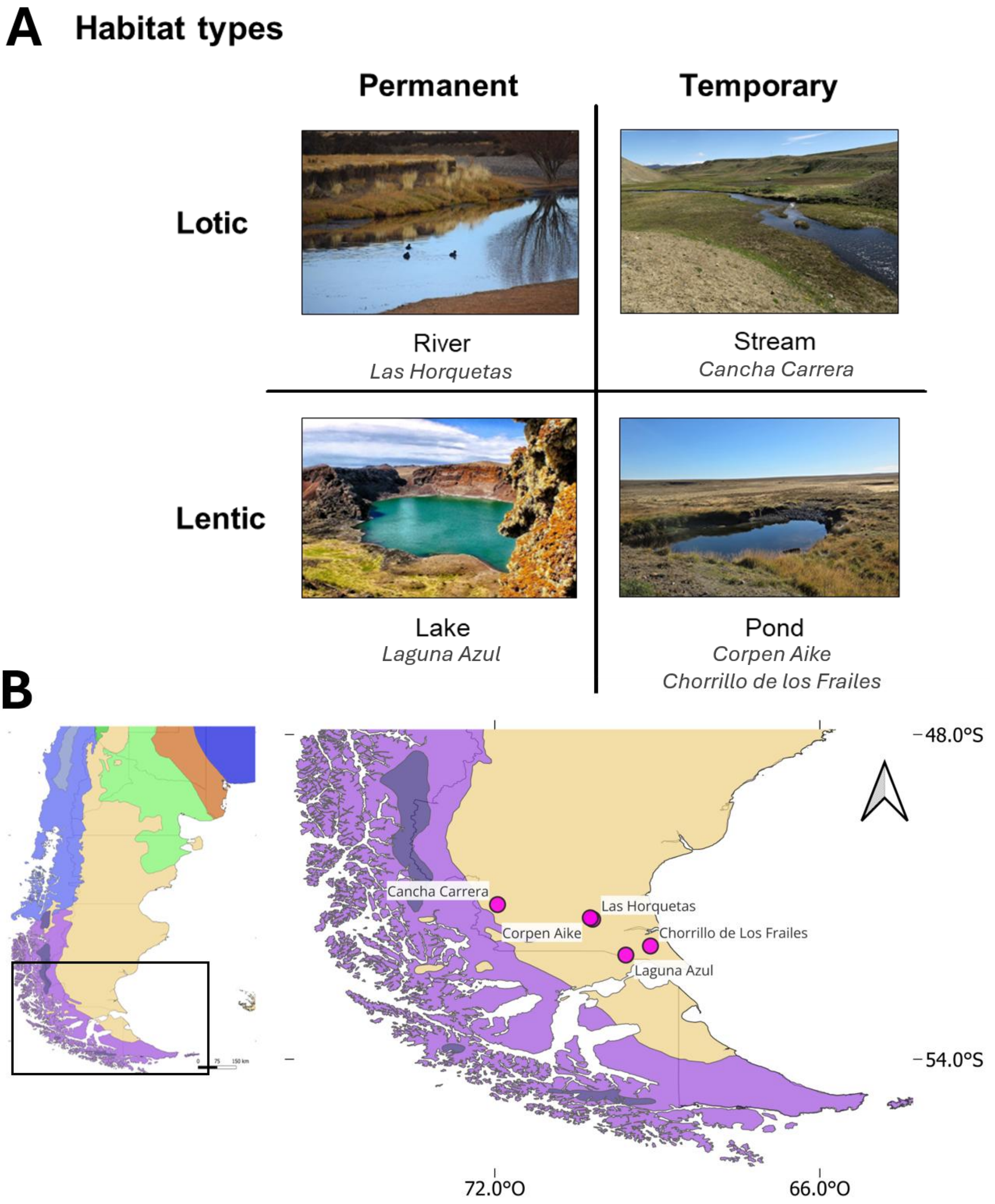

**Figure S2.** Experimental breeding design. Wild-caught individuals ($G_0$; 80–100 snails per population) from two lentic populations of *Pectinidens diaphanus* exhibiting contrasting shell morphologies—a permanent lake population and a temporary pond population—were transferred to the laboratory and maintained under standardized conditions (16 °C, 16 h light : 8 h dark photoperiod, food provided *ad libitum*). For each population, $G_0$ individuals were split in two aquaria (A and B), where they reproduced through mass mating and produced $G_1$ egg clutches. After hatching, juvenile snails from aquaria A and B were intermixed within each population and redistributed to establish experimental populations. $G_1$ individuals were reared to adulthood and allowed to reproduce through mass mating, generating $G_2$ offspring. The same procedure was repeated across generations without interpopulation crosses. Shell morphology was subsequently compared between wild-caught $G_0$ individuals and laboratory-reared $G_2$ descendants to assess the persistence of shell-shape differences under common-garden conditions.

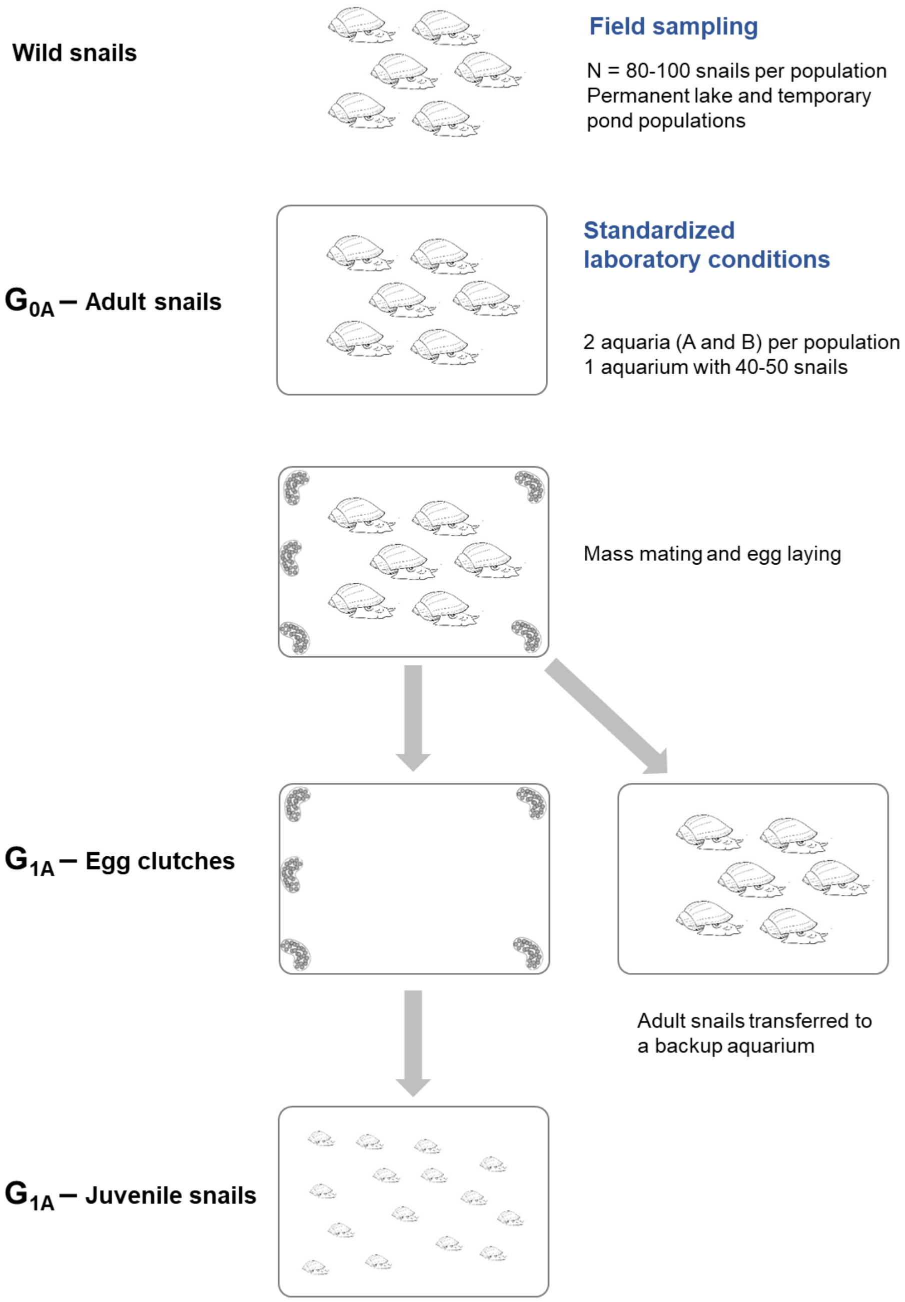

**Figure S2.** *Continued…*

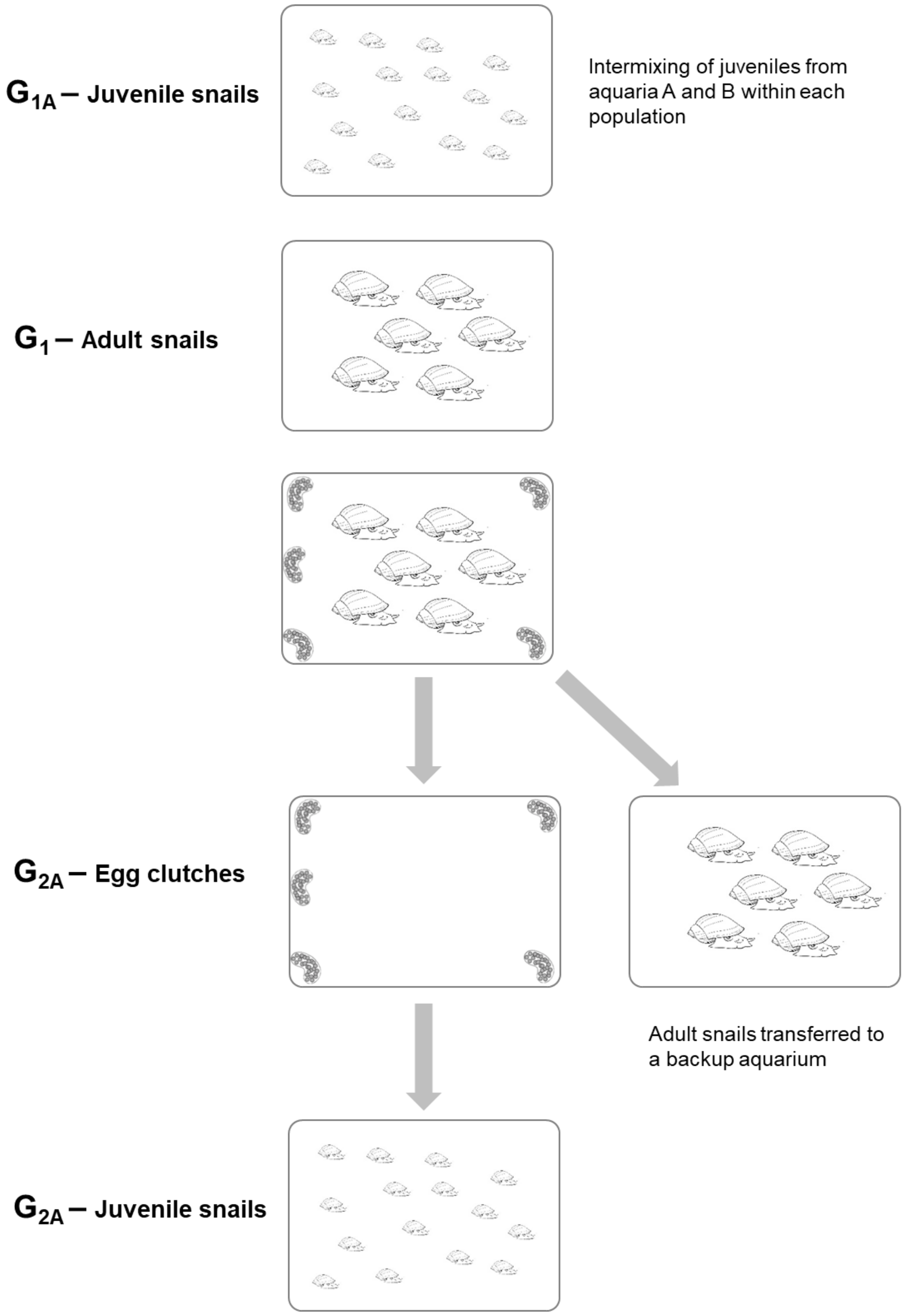

**Figure S3.** Dorsal view of the anatomy of the reproductive system of an adult snail *Pectinidens diaphanus* collected from the lake Laguna Azul in Santa Cruz Province, Argentina. The individual was dissected under a stereoscopic microscope and drawing of the reproductive system was made using a camera lucida attachment. The pictures obtained were digitalized with an 'Apple Scanner'. ag = albumen gland; ng = nidamental gland; ot = ovotestis; pr = prostate; pp = preputium; ps = penis sheath; sd = spermiduct; sp = spermatheca; sv = seminal vesicle; va = vagina; vd = vas deferens. Bar indicates 1 mm.

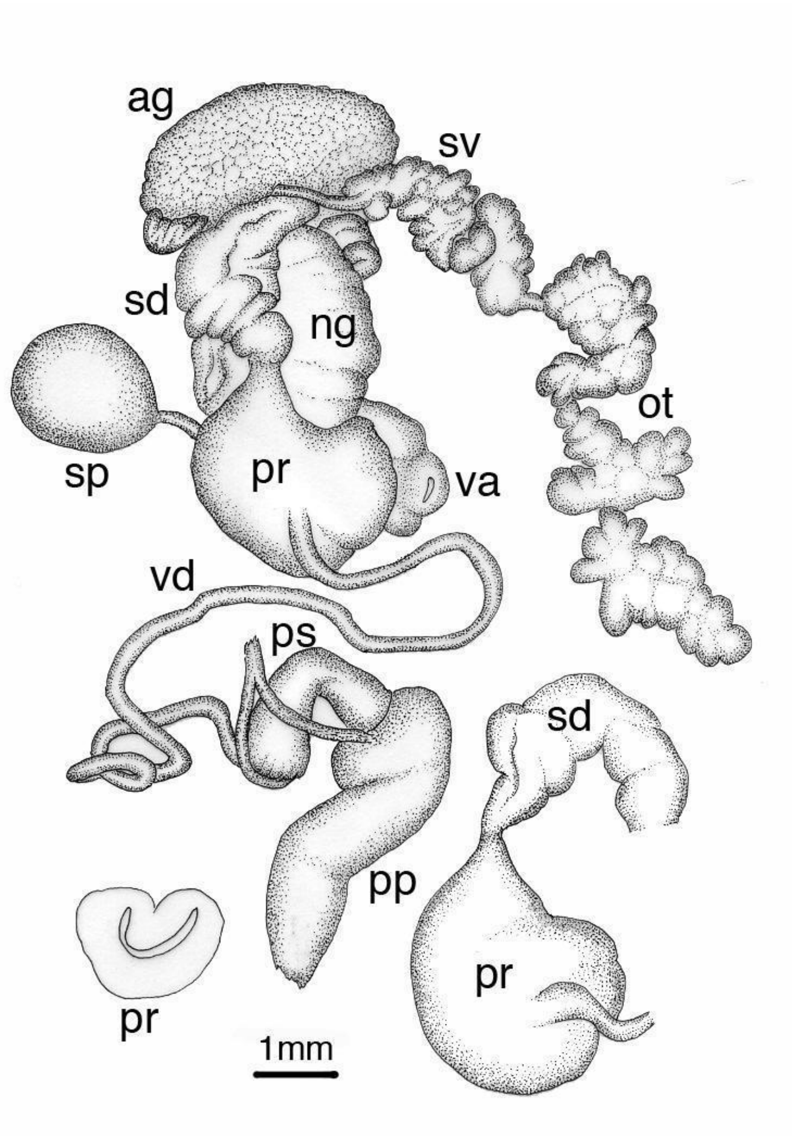

**Figure S4.** TCS haplotype network based on COI sequences (473–627 pb) of *Pectinidens diaphanus* and the outgroup *Ladislavella edoles*. Each circle represents a unique haplotype, and its size is proportional to the number of individuals sharing that haplotype. Each hatch mark represents a single nucleotide substitution. Small black circles represent hypothetical (unsampled) haplotypes.

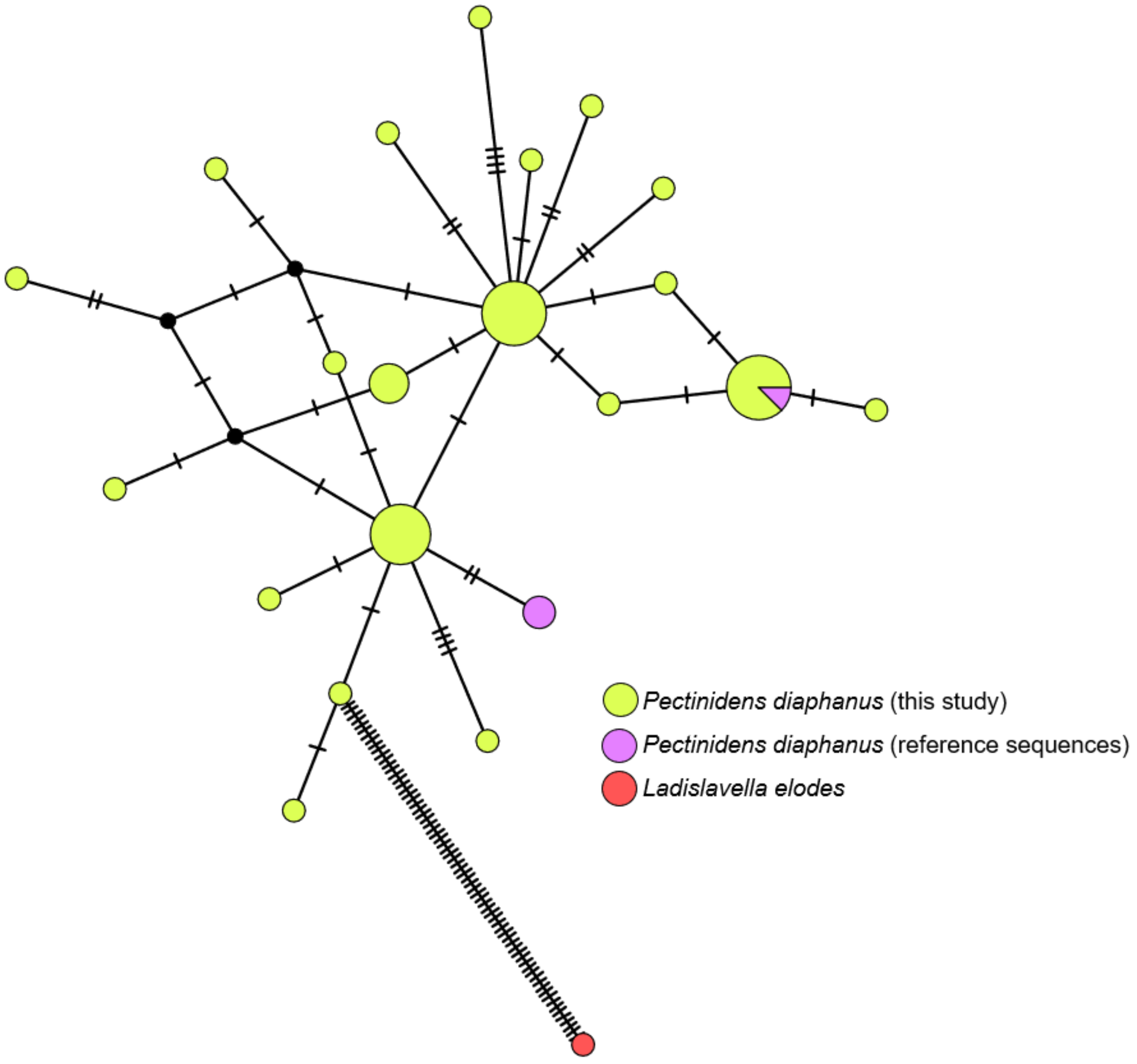
